# Variability within rare cell states enables multiple paths towards drug resistance

**DOI:** 10.1101/2020.03.18.996660

**Authors:** Benjamin L. Emert, Christopher Cote, Eduardo A. Torre, Ian P. Dardani, Connie L. Jiang, Naveen Jain, Sydney M. Shaffer, Arjun Raj

## Abstract

Molecular differences between individual cells can lead to dramatic differences in cell fate, such as death versus survival of cancer cells upon drug treatment. These originating differences remain largely hidden due to difficulties in determining precisely what variable molecular features lead to which cellular fates. Thus, we developed Rewind, a methodology that combines genetic barcoding with RNA FISH to directly capture rare cells that give rise to cellular behaviors of interest. Applied to BRAF^V600E^ melanoma, we trace drug-resistant cell fates back to single-cell gene expression differences in their drug-naive precursors (initial frequency of ∼1:1000-1:10,000 cells) and relative persistence of MAP-kinase signaling soon after drug treatment. Within this rare subpopulation, we uncover a rich substructure in which molecular differences between several distinct subpopulations predict future differences in phenotypic behavior, such as proliferative capacity of distinct resistant clones following drug treatment. Similarly, we show that treatments that modify the frequency of resistance can allow otherwise non-resistant cells in the drug-naive population to become resistant, and that these new populations are marked by the variable expression of distinct genes. Together, our results reveal the presence of hidden, rare-cell variability that can underlie a range of latent phenotypic outcomes upon drug exposure.

## Introduction

Individual cells—even those of ostensibly the same cell type—can differ from each other in a number of ways. Some of these differences can result in a “primed” cellular state that can, in a particular context, ultimately lead to biologically distinct behaviors ^1,2^. This cellular priming underlies a number of important single-cell phenomena. For instance, when anti-cancer therapeutics are applied to clonally derived cancer cells, most of the cells die; however, a small number of cells survive and proliferate, and these cells drive therapy resistance ^3–6^. Yet, while this phenomenon suggests the existence of rare, primed cells in the initial population, it remains unclear what distinguishes these cells at the molecular level from the rest of the population.

We and others have shown that rare cells within an isogenic population can exhibit fluctuations in expression of several genes simultaneously, which predict rare-cell phenotypes and persist through multiple cell divisions ^3,7^. What remains largely unknown, outside of a few cases ^6,8,9^, is precisely how this variability maps to distinct cellular outcomes following a treatment. As a result, several questions remain unanswered. Is molecular variability in the initial state of cells inconsequential because all cells ultimately funnel into the same cell fate? Can different cell fates arise from otherwise indistinguishable initial molecular states? Or can most differences in ultimate fate be traced back to measurable differences in the initial states of cells? What is the structure of this initial variability? These questions remain largely unanswered because of our limited ability to longitudinally track and profile cells (especially rare ones) from initial state to final fate. Longitudinal profiling by time-lapse microscopy is generally limited in its ability to interrogate large numbers of molecular features simultaneously ^8,10^. Barcoding, in which cells are labeled by unique and sometimes mutable nucleic acid sequences ^11–16^, allows one to track and profile single cells by sequencing or imaging based readouts ^17–20^. However, a key challenge for both of these methodologies is the detection of rare cells (1:1000 or even more rare), for which neither time-lapse nor single-cell RNA sequencing is particularly effective (new techniques aim to circumvent these limitations ^21–24^). Yet, many biological phenomena, such as therapy resistance in cancer cells, occur in subpopulations that are at least that rare.

Here, we explicitly connect drug-resistant cell fates in melanoma to specific molecular features in rare subsets of cells in the drug-naive population. These connections revealed a rich mapping between previously hidden single-cell variability and a number of latent cellular behaviors. Our results suggest the existence of a large number of rare subpopulations within seemingly homogenous cells, each with potentially distinct biological behaviors, and set out a path to discover biologically consequential axes of variability.

## Results

### Rewind enables retrospective identification of rare cell populations

Therapy resistance in cancer provides an excellent system in which to map out the connections between rare cell states and fates. In this context, fates refer to cells that proliferate when treated with targeted therapies, and the states are the molecular profiles of drug-naive cells that will ultimately lead to these resistant fates. These variable profiles can appear even in clonally derived lines and have a non-genetic basis ^3-6^. We here have focused on BRAF^V600E^-mutated melanoma, in which we have previously demonstrated that there is a rare, transient subpopulation composed of cells (∼1:2000) that are “primed” to survive treatment to the targeted therapy vemurafenib ^7,25^. These rare, primed cells often express higher levels of certain receptor tyrosine kinases (such as *EGFR, NGFR* and *AXL*) and lower levels of melanocyte-determining transcription factors (*SOX10* and *MITF*) than the rest of the cells in the population. However, these markers are highly imperfect, with many positive cells being non-resistant and many negative cells being resistant, leaving open the question as to what markers specifically mark the primed state.

The primary technical challenge for studying rare cell processes like drug resistance is the rarity of the cells of interest. Current techniques for retrospective identification require profiling of the entire initial population and then post-facto determining which profiles correspond to cells of interest ^17,18^. We developed an alternative methodology, dubbed Rewind, to retrospectively isolate or identify rare cell populations of interest for downstream characterization. Rewind works by using a lentiviral library of transcribed barcodes, in which the barcode sequence is incorporated into the 3’ untranslated region of green fluorescent protein (GFP) mRNA (Fig. 1a and Supplementary Fig. 1a). After labeling cells with these barcodes, we allowed the cells to divide for a few divisions and then separated the population into two equal groups (“twins”) such that most barcoded lineages (>90%) were present in each group (see Methods for discussion and empirical simulations). One group we fix in time as a “Carbon Copy” of the cells in their initial state, and to the other, we apply the treatment to see which cells undergo the rare behavior of interest (e.g., becoming resistant to drug). After selecting the cells that undergo the rare behavior, we sequence their DNA to identify their barcodes, and then we use those barcodes to identify their “twins” in the Carbon Copy by fluorescently labelling the RNA transcribed from those specific barcodes using RNA in-situ-hybridization techniques (Supplementary Fig. 1b,c,f,h). We verified that the barcode library was sufficiently diverse to label 100,000s of cells with over 99% receiving unique barcodes, thus minimizing spurious identification (see Methods and Supplementary Fig. 2 for experimental details and calculations). Once isolated, we can molecularly profile the Carbon Copy twins to determine what is different about their initial state that led to their distinct fate. Altogether, the Rewind methodology enables retrospectively uncovering primed cell states that lead to rare cell behaviors.

**Figure 1:**
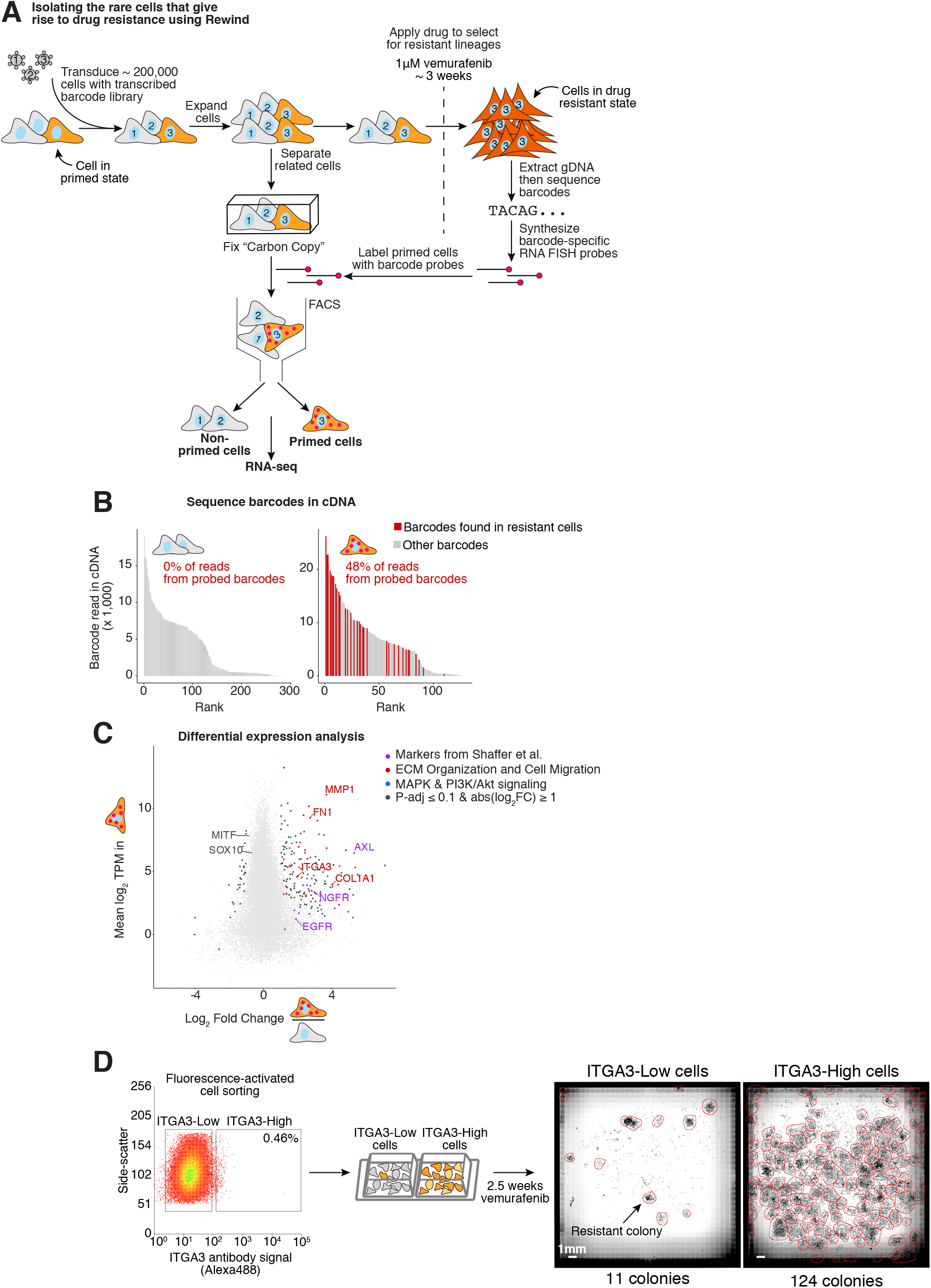
Rewind identifies rare cell states giving rise to vemurafenib resistant colonies. **A**. Schematic of Rewind approach for isolating the initial primed WM989 A6-G3 melanoma cells that ultimately give rise to vemurafenib resistant colonies. For the experiment shown, we transduced ∼ 200,000 WM989 A6-G3 cells at an MOI ∼ 1.0 with our Rewind barcode library. After 11 days (∼4 population doublings) we divided the culture in two, fixing half in suspension as a Carbon Copy and treating the other half with 1 μM vemurafenib to select for resistant cells. After 3 weeks in vemurafenib, we extracted genomic DNA from the resistant cells that remain and identified their Rewind barcodes by targeted sequencing. We then designed RNA FISH probes targeting 60 of these barcodes and used these probes to specifically label cells primed to become resistant from our Carbon Copy. We then sorted these cells out from the population, extracted cellular RNA and performed RNA sequencing. **B**. To assess the sensitivity and specificity of the Rewind experiment in A, we performed targeted sequencing to identify barcodes from cDNA generated during RNA-seq library preparation. Bar graphs show the abundance (y-axis) and rank (x-axis) of each sequenced barcode (≥ 5 normalized reads). Red bars correspond to barcodes targeted by our probe set and gray bars correspond to “off-target” barcode sequences. Inset shows the percent of barcode sequencing reads that match a probe-targeted barcode. These data correspond to 1 of 2 replicates. **C**. We performed differential expression analysis using DESeq2 of primed vs. non-primed sorted cells. Shown is the mean expression level (log_2_(transcripts per million)) for protein coding genes in primed cells (y-axis) and log_2_ fold change in expression estimated using DESeq2 (x-axis) compared to non-primed cells. Colors indicate differentially expressed genes related to ECM Organization and Cell Migration (red), MAPK and PI3K/Akt signaling pathways (blue) and previously identified resistance markers ^6^ (purple). Genes were assigned to categories based on a consensus of KEGG pathway and GO enrichment analyses (see Methods for details). **D**. We selected the most differentially expressed, cell surface ECM-related gene (*ITGA3*) to validate as a predictive marker of vemurafenib resistance in WM989 A6-G3. After staining cells with a fluorescently labelled antibody targeting ITGA3, we sorted the brightest 0.5% (ITGA3-High) and remaining (ITGA3-Low) populations, then treated both with 1 μM vemurafenib. After approximately 18 days, we fixed the cells, stained nuclei with DAPI then imaged the entire wells to quantify the number of resistant colonies and cells. The data correspond to 1 of 3 biological replicates (see Supplementary Fig. 4 for additional replicates).

A critical feature of these rare primed cell states is that they are transient, meaning that cells can fluctuate both into and out of the primed state ^3,6^. An important biological question that is relevant to the ability of Rewind to profile primed cells is whether these cells maintain (“remember”) their primed state through several cell divisions. (Memory would be required for the profile of cells isolated from the Carbon Copy to reflect those of their twins that received treatment with vemurafenib.) To empirically test for the existence of such memory, we let a barcoded WM989 A6-G3 culture double 4-5 times, split the culture in two, and then separately treated both halves of the population with vemurafenib. We found a large overlap in the barcodes between the two halves, demonstrating that the primed state is maintained for several divisions and that there is sufficient memory in the system for Rewind to effectively profile the primed state (Supplementary Fig. 3).

### Tracing vemurafenib-resistant melanoma cells back to their rare, drug-naive precursors for gene expression profiling

We then applied the Rewind approach to isolate the rare WM989 A6-G3 cells primed for vemurafenib resistance by FACS, after which we profiled these primed drug-naive cells by RNA sequencing (Fig. 1 and Supplementary Fig. 4a). Upon sequencing barcodes from cDNA, we found that ∼48% of reads in the sorted primed subpopulation contained probe-targeted barcodes matching those identified in vemurafenib resistant colonies (vs. 0% in the non-primed subpopulation), reflecting an estimated ∼1,600-fold enrichment over the baseline frequency of these barcodes in the Carbon Copy (∼0.03%; Fig. 1B). (We suspect the proportion of on-target cells isolated here is lower than in our pilot experiments (Supplementary Fig. 1b,c) due to the lower prevalence of the targeted cells.) Having confirmed that FACS enriched for primed cells, we then looked for differentially expressed genes compared to non-primed cells. Consistent with previous research from our lab and others, we found that primed cells sorted from the Carbon Copy expressed greater than 2-fold higher levels of the receptor tyrosine kinases *AXL, EGFR* and *NGFR* as well as lower levels of the melanocyte transcription factors *SOX10* and *MITF* (Supplementary Fig. 4c) ^6,26^. Beyond these known markers, the transcriptome profile provided by Rewind enabled us to identify nearly 200 new marker genes whose expression was significantly altered in primed cells. Among these genes, we found a significant enrichment for genes associated with cell adhesion, extracellular matrix (ECM) organization and cell migration (Fig. 1c, Supplementary Fig. 4d and Supplementary Table 6). Longitudinal tracking of primed cells revealed that the expression of most priming marker genes either stayed the same or increased during the acquisition of stable resistance over 3 weeks in vemurafenib treatment, while an additional ∼2,800 genes showed a greater than 2-fold change in expression during this period (Supplementary Fig. 5). Thus, most of the genes that are upregulated in resistant cells are not the genes whose expression marks the primed state, thus motivating the use of Rewind to identify these markers.

Many of these markers have not previously been implicated in cellular priming for vemurafenib resistance and hence represent potentially novel single-cell biomarkers of resistance. An example was *ITGA3*, which was the most differentially expressed cell surface marker identified by Rewind. To verify that it marked primed cells, we prospectively sorted drug-naive WM989 A6-G3 cells expressing high levels of *ITGA3*. These cells gave rise to 10-fold more resistant colonies upon exposure to vemurafenib, confirming that ITGA3 is a marker (Fig. 1d and Supplementary Fig. 4e-h). We also used Rewind to identify markers in another melanoma line, WM983b E9-C6, in which markers of the cells primed for resistance were unknown, revealing and validating that *AXL* was a marker (Supplementary Fig. 6). Together, these results demonstrate that there are large sets of genes that exhibit rare-cell fluctuations that can lead to drug resistance.

### Individual primed cells are marked by coordinated expression of multiple resistance markers prior to vemurafenib treatment

Yet, while isolating rare cells that express high levels of these markers enriched for cells that could become drug resistant, we also observed that the majority of cells that expressed any one marker still died when faced with drug. Thus, there was no one factor whose expression precisely marked the cells that were primed for drug resistance. These facts suggest that the cellular fluctuations that lead to a cell becoming primed for drug resistance may be complex, and potentially marked by the fluctuations of several genes in tandem. Indeed, our lack of knowledge of the precise nature of the mapping between fluctuations and outcomes leaves open a rich set of possibilities. In principle, rare-cell fluctuations of genes associated with a particular behavior need not be independent of each other, but may take on many correlation structures and sub-structures, with sets of genes potentially co-fluctuating or anti-fluctuating to demarcate specific subpopulations within the overall rare-cell population. A parallel question is whether these different subpopulations all funnel to the same drug-resistant outcome: it is possible that these new axes of variability may represent fluctuations that lead primed cells to adopt phenotypically distinct cellular fates after, say, the addition of drug. Rewind allowed us to look for these new sub-populations.

We first attempted to resolve the question of why most cells that expressed any one particular marker actually did not become resistant to drug. We hypothesized that simultaneous co-expression of multiple markers may more accurately and specifically identify the exact cells that are primed to be resistant. To look for evidence of such structured fluctuations, we used Rewind in combination with RNA imaging to transcriptionally profile primed cells with single-molecule resolution (Fig. 2a,b). In this manner, we located 162 primed cells in situ within a total of ∼750,000 cells scanned in our Carbon Copy, which we then probed for expression of 9 genes by single-molecule RNA FISH (Methods). These cells showed substantially higher expression of *AXL, EGFR, NGFR, WNT5A, ITGA3, MMP1*, and *FN1* and lower expression of *SOX10* and *MITF* than randomly selected cells, consistent with our earlier results from RNA-seq (Fig. 2c,d). Overall differences in expression capacity were unlikely to explain the increased expression of marker genes in primed cells (Supplementary Fig. 4b, Supplementary Fig. 4h and Supplementary Fig. 7e). Moreover, cells primed for resistance were far more likely to co-express any pair of markers (Odds Ratios ranging from ∼1.5 to ≥58; Supplementary Fig. 7), and ∼87% percent of cells expressed high levels of ≥4 of 7 marker genes simultaneously, in stark contrast to cells not expressing resistant barcodes (Fig. 2e and Supplementary Fig. 7). This apparent coordination suggests that the cell-to-cell differences that lead to distinct cell fates following drug treatment are a consequence of the coordinated fluctuations of several factors simultaneously, as opposed to sporadic fluctuations of individual genes ^7^.

**Figure 2:**
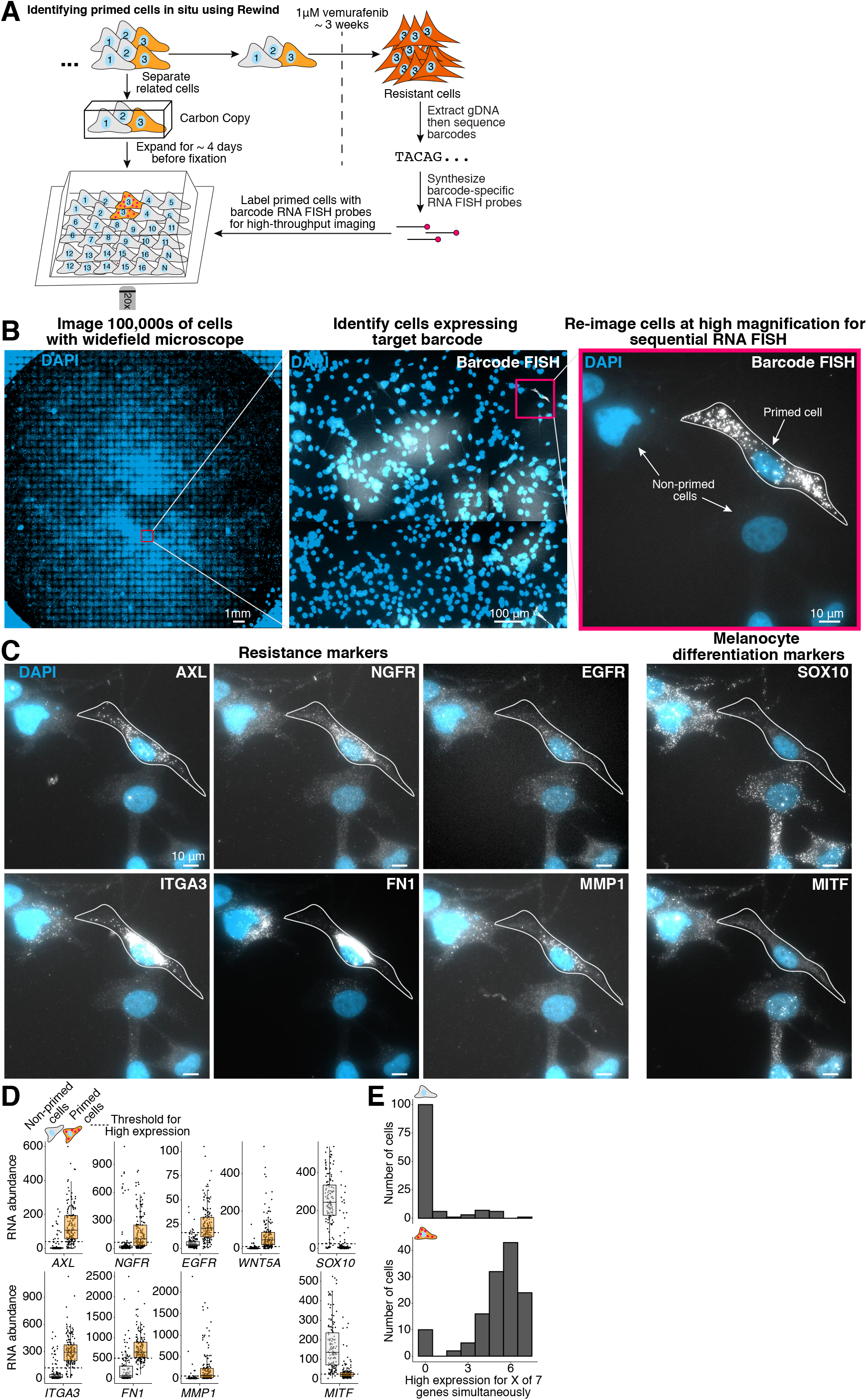
A coordinated primed cell state characterized by high expression of multiple markers gives rise to vemurafenib resistance in WM989 A6-G3. **A**. We performed Rewind with image-based profiling to identify WM989 A6-G3 cells primed to become vemurafenib resistant in situ and measure gene expression in individual cells using single-molecule RNA FISH. We expanded barcoded cells for ∼4 population doublings before dividing the cells into the Carbon Copy or the drug-treated half. **B-C**. To identify the rare primed cells, we first imaged Carbon Copies at 20X magnification and identified primed cells labeled with our barcode RNA FISH probes using a combination of automated image analysis and manual image review. Once identified, we returned to these cells (n = 162) for re-imaging at high magnification (60X) and quantification of marker gene expression using single-molecule RNA FISH. We additionally imaged multiple randomly selected positions in each well to quantify marker gene expression in “non-primed” cells (n = 135). **D**. Quantification of single-cell gene expression in primed and non-primed cell subpopulations. Each point corresponds to an individual cell. We set thresholds for high marker expression based on the observed expression distribution in non-primed cells (see Methods and Supplementary Fig. 7 for details). **E**. Frequency of cells expressing high levels (beyond the thresholds shown in D.) of 1, 2,…7 markers (out of a total of 7 measured) simultaneously in primed and non-primed cell populations. The number of cells from each subpopulation with data for all 7 markers are indicated above each histogram. These data correspond to 1 of 2 biological replicates (see Supplementary Fig. 7 for the additional replicate).

### Primed melanoma cells are marked by higher levels of phosphorylated ERK shortly after, but not prior to, vemurafenib treatment

A possible mechanism for how these primed cells survive drug treatment is that the observed increases in expression of multiple receptor tyrosine kinases and their cognate ligands lead to differences in MAPK pathway activation. To address this hypothesis we measured dual phosphorylated ERK (pERK) levels in primed and non-primed cells by immunofluorescence (Fig. 3 and Supplementary Fig. 8). We found similar levels of pERK in primed and non-primed cells in Carbon Copies fixed before vemurafenib treatment. However, in Carbon Copies that underwent vemurafenib treatment for 24 hours, we found that primed cells had residual levels of pERK that were on average 40% higher than the rest of the population, with some primed cells having levels nearly 5-fold higher than non-primed cells (within the range of untreated cells). We also observed that within individual clusters of closely related primed cells, not all cells contained higher levels of pERK, which may reflect pulsatile changes in pERK as documented elsewhere (Supplementary Fig. 8e) ^27^. In contrast, single-cell levels of total ERK levels were modestly lower in primed cells compared to non-primed cells, both before and after vemurafenib treatment (Fig. 3d and Supplementary Fig. 8b). These results suggest that primed cells are able to maintain residual MAPK signaling following vemurafenib treatment that may allow them to continue proliferating in the face of drug.

**Figure 3:**
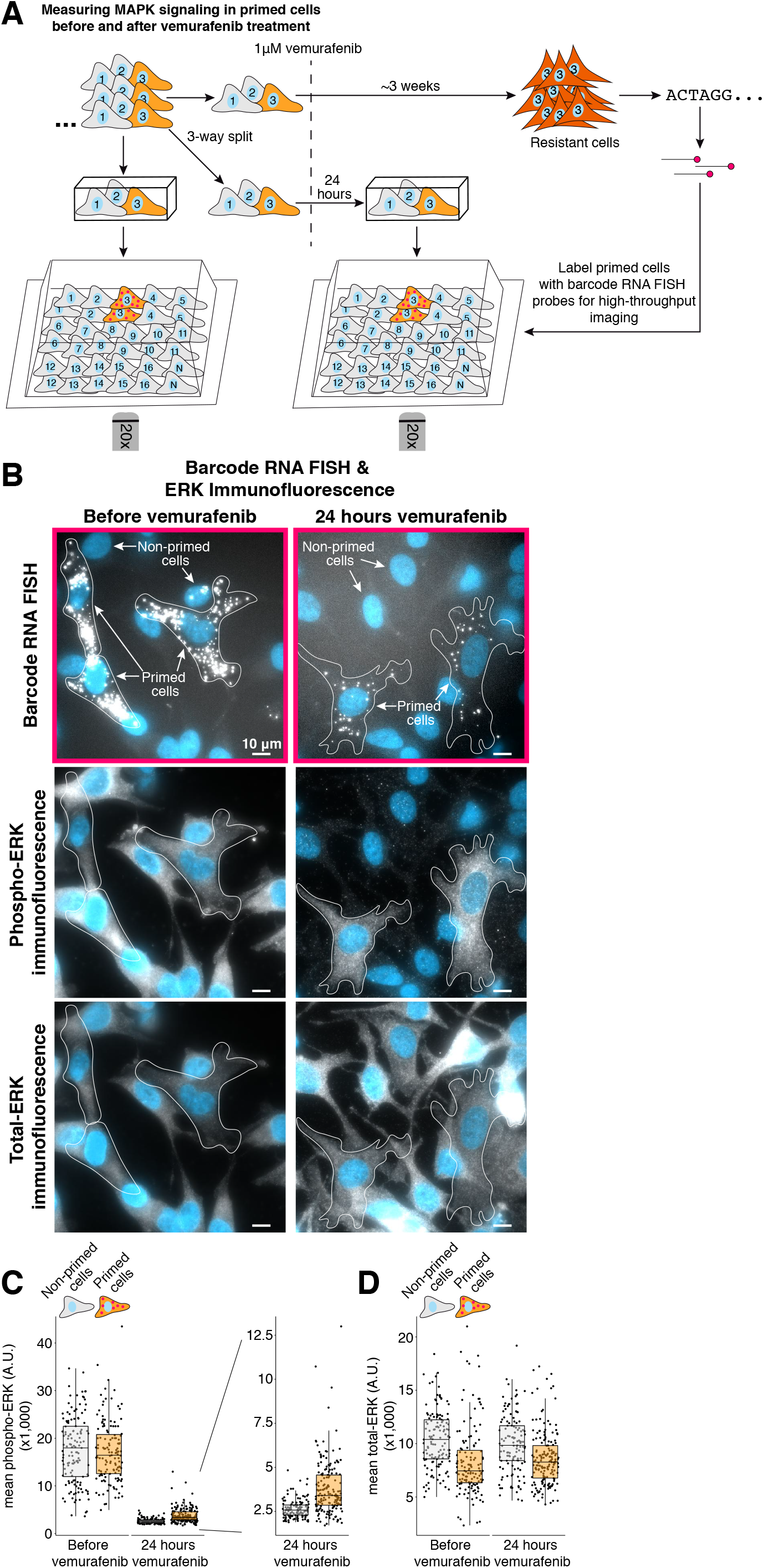
Resistance to vemurafenib is associated with single-cell variability in phosphorylated ERK levels 24 hours after treatment but not prior to treatment. **A**. We used Rewind to quantify dual-phospho ERK (p44/p42, pERK) and total ERK levels in primed cells before and 24 hours after vemurafenib treatment. To quantify ERK levels over time, we expanded barcoded cells for ∼4 population doublings then plated two Carbon Copies and fixed one 24 hours after vemurafenib treatment and the other prior to treatment. As before, we used barcode RNA FISH probes to identify primed cells in both Carbon Copies then measured single-cell levels of total ERK and pERK by immunofluorescence (n = 135 cells without vemurafenib treatment and n = 173 cells with vemurafenib treatment). We additionally imaged multiple randomly selected positions in each well to quantify total ERK and pERK in non-primed cells (n = 133 cells without vemurafenib treatment and n = 125 cells with vemurafenib treatment). **B**. Barcode RNA FISH and ERK immunofluorescence images of primed cells identified in Carbon Copies fixed before vemurafenib treatment (left) and 24 hours after treatment (right). **C-D**. Quantification of average pERK and average total ERK immunofluorescence intensity in primed cells and non-primed cells. Each point corresponds to an individual cell. These data correspond to 1 of 2 biological replicates (see Supplementary Fig. 8 for the additional replicate).

### Distinct drug-resistant fates can be traced back to molecular differences within the primed subpopulation

While these results showed an overall coordination between the different marker genes in primed cells, there were considerable differences in the degree of co-expression between these marker genes in single cells (Supplementary Fig. 7c,d,h,i). These differences suggest the possibility that the expression of specific subsets of genes may delineate specific subpopulations within the overall rare primed population that could in principle have different fates. Evidence for different fates comes from inspection: it was visually clear that different colonies of vemurafenib-resistant cells can show dramatic differences in basic properties like the number of cells in the colony. We wondered whether tracing back these differences in fate with Rewind could reveal the molecular profiles that distinguish subsets of the initial primed cell subpopulation with distinct potential. We applied Rewind in the WM989 A6-G3 cell line as before, but used the number of barcode reads in the resistant population as a proxy for the number of resistant cells carrying a given barcode. We then designed RNA FISH probes that distinguished 30 of the most abundant barcodes (i.e., “highly resistant”, meaning many resistant cells) from 30 barcodes in the next tier of abundance (i.e., “less resistant”). We used these probes to identify their twin cells in a Carbon Copy fixed prior to vemurafenib treatment (Fig. 4 and Supplementary Fig. 9 for probe set validation).

**Figure 4:**
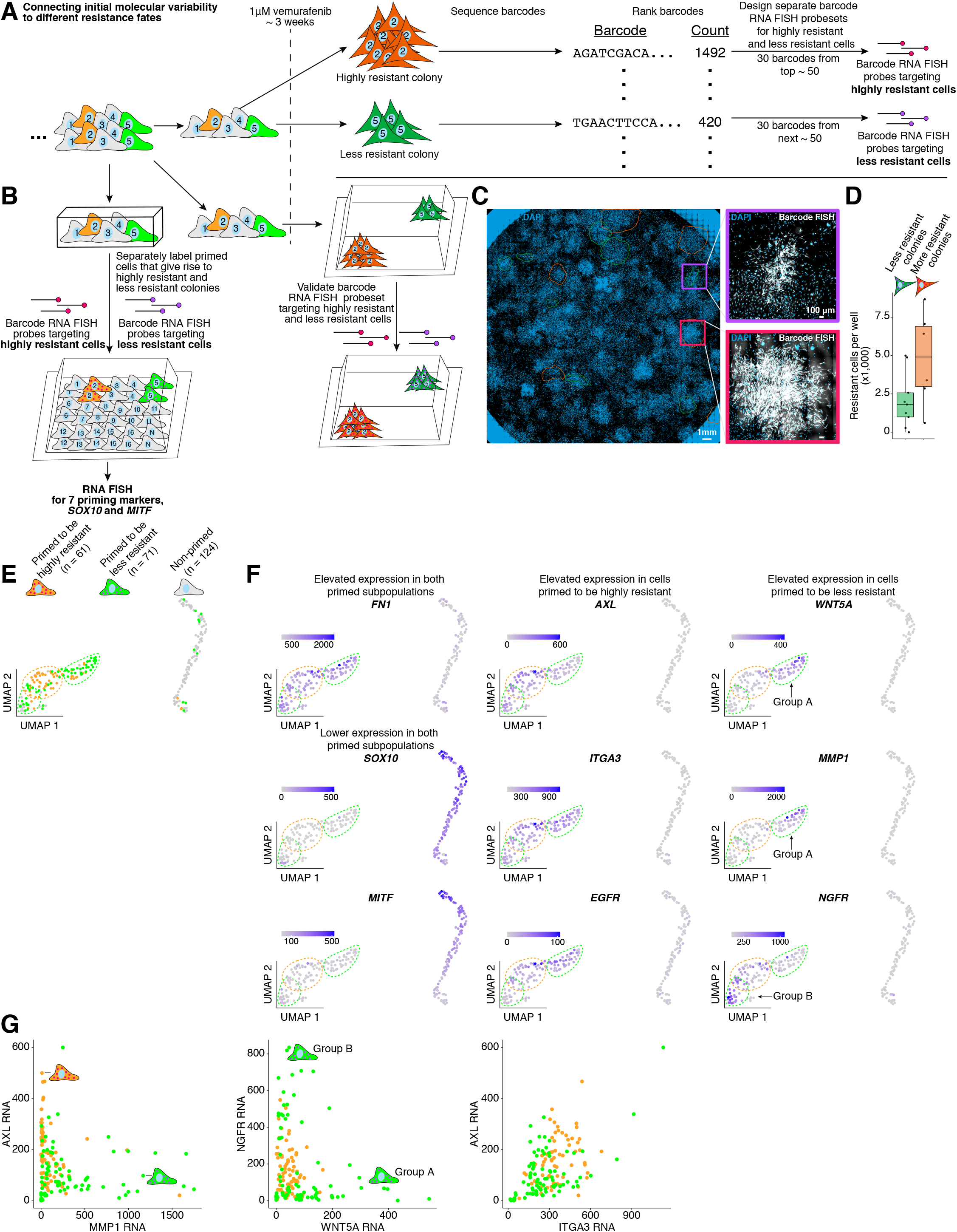
Variation in gene expression among primed cells is associated with differences in resistant cell fate. **A**. We performed Rewind in WM989 A6-G3 cells and identified barcode sequences enriched in resistant colonies following vemurafenib treatment. We ranked these barcodes by abundance as a proxy for ranking the number of resistant cells carrying each specific barcode. We then designed separate RNA FISH probe sets targeting barcodes from the ∼ 50 most abundant resistant clones (“highly resistant cells”) and barcodes targeting the next ∼ 50 resistant clones (“less resistant cells”). Each probe set contained probes targeting 30 distinct barcodes. **B**. We used these separate probe sets to identify corresponding primed cells in the Carbon Copy fixed prior to vemurafenib treatment then performed sequential rounds of RNA FISH to measure single-cell expression of 9 genes. We additionally imaged multiple randomly selected positions to quantify gene expression in non-primed cells. These data are the same as used in Fig. 2, here analyzed using information on which probe set labeled each cell. **C-D**. To check whether the separate probe sets label barcode RNA corresponding to distinct resistant fates, we labelled resistant colonies derived from the same population of cells, then quantified the number of resistant cells labelled with each probe set. The number of colonies labeled with each probe set and the average number of cells per colony are shown in Supplementary Fig. 9. These data correspond to 1 biological replicate. **E**. Using the RNA FISH data from the Carbon Copy in B., we applied the UMAP algorithm to the first 5 principal components to visualize differences in gene expression between primed cells (n = 132) and non-primed cells (n = 124). We then colored each cell by its predicted fate based on its barcode. To orient the reader, we circled the largest group of primed cells that give rise to highly resistant colonies in orange, and the two separate groups of primed cells that give rise to less resistant colonies in green. **F**. Maintaining the organization provided by UMAP, we colored each cell by its expression of each of the 9 genes measured. As noted in the text, ≥98% of primed cells had levels of FN1 RNA that were 3-fold higher than the median observed in non-primed cells, and ≥80% of primed cells had levels of SOX10 and MITF RNA that were ≤ ⅓ the median levels observed in non-primed cells. **G**. Scatterplots show the single-cell expression for pairs of markers that distinguished the groupings shown in D.

To find transcriptional profiles that predict whether cells are primed to become either highly resistant or less resistant, we measured transcript abundances in individual primed cells by RNA FISH for 9 genes, including 7 priming markers, *MITF* and *SOX10*. We used the dimensional reduction technique UMAP to visualize differences between cells based on expression levels. We then marked individual cells in this visualization based on their ultimate fate as determined by the barcode RNA FISH signal (primed to become highly vs. less resistant vs. non-primed). We found that non-primed cells clearly separated from all the primed cells, and that within the primed cells, the highly resistant primed cells grouped together, while the less resistant cells formed two distinct groups (Fig. 4e,f). These groupings were also apparent in hierarchical clustering of the single-cell gene expression data, with cluster assignment of each cell roughly corresponding to its resistance fate, suggesting a clear distinction between the groupings (Supplementary Fig. 10c,e).

We then asked how expression levels of particular genes corresponded to these groupings. As expected, most (>80%) of the primed cells had markedly decreased levels of both *SOX10* and *MITF* (Fig. 4f, Supplementary Fig. 7, and Supplementary Fig. 10c). We also found that almost all primed cells had increased levels of *FN1* (>98%), thus suggesting that *FN1* is a “pan” marker of cells primed for vemurafenib resistance (Fig. 4F, Supp. Fig. 7, and Supplementary Fig. 10c). Co-expression of *AXL, ITGA3*, and *EGFR* marked cells primed to become highly resistant, but individually these genes were also expressed in subsets of cells primed to become less resistant (Fig. 4f and Supplementary Fig. 10c). These subsets could also be distinguished by expression of *WNT5A, MMP1, NGFR with* one group (group A) expressing the highest levels of *WNT5A* and *MMP1* and the other (group B) expressing the highest levels of *NGFR* (*NGFR* also had intermediate levels of expression in the cells primed to be highly resistant; Fig. 4f). In addition, quantitative comparison of expression levels between pairs of markers showed that the expression of, for example, *AXL* vs. *MMP1* fell along two separate axes of variability (Fig. 4g). Together, these analyses suggest that multiple classes of primed cells with different expression patterns give rise to resistant colonies with different phenotypes.

Although our labeling scheme did not discriminate between different primed cells that ended up with the same fate, in these imaging data, we were able to use spatial proximity of barcode-positive cells to infer that neighboring barcode-positive cells were likely derived from the same initial cell and therefore belong to a unique subclone (Supplementary Fig. 10b). We could then use the single-cell gene expression levels to further determine which primed cell class these cells belonged to, and ask whether there were any signs of switching between primed cell classes (including reversion to the non-primed state) (Supplementary Fig. 10). In nearly half of the subclones (11 out of 24), all cells fell into a single primed-cell class. Moreover, for most (7 out of 13 remaining) subclones containing a mix of cell states, only 1 cell within the subclone was classified as a separate class (Supplementary Fig. 10d right). These data suggest that primed cells can transition between states, and these transitions occur on a relatively slow time-scale (potentially once per 4 days or ∼2-3 cell divisions; slow compared to most expression fluctuations), consistent with recent work quantifying the transcriptional memory of several primed-cell marker genes ^7^.

### DOT1L inhibition enables a distinct primed subpopulation of melanoma cells to become vemurafenib resistant

These results show that primed cells consist of a complex set of subpopulations that can map to a variety of cell fates. A critical point is that the mapping and hence subpopulations were revealed by the addition of a particular drug. It is possible that there are additional subpopulations present in cells that would normally not survive drug treatment. Further, it may be that the molecular differences that characterize these subpopulations could allow otherwise drug-susceptible cells to become primed for drug resistance in different conditions. Evidence for such a possibility comes from the existence of factors that, when perturbed in drug-naive cells, can reduce or increase the frequency of resistant colony formation, implying an increase or decrease in the number of primed cells within the population ^25^. Amongst these is DOT1L, a H3K79 methylase whose inhibition leads to a 3-fold increase in the number of resistant colonies that form upon addition of vemurafenib ^25^. While DOT1L inhibition removes some type of barrier that allows more cells to be primed, this barrier is not removed in all cells because not all cells are able to form resistant colonies. Thus, an important question is what distinguishes the small subset of the cells that become primed for resistance upon DOT1L inhibition from the majority of cells that remain non-resistant to drug. (Barcoding analysis revealed that DOT1L inhibition indeed permits a new subset of cells to enter a primed state rather than affecting proliferation or reversion of primed cells; Supplementary Fig. 11.)

Using Rewind, we sought to reveal the molecular profile specific to the subpopulation of cells that required DOT1L inhibition to survive vemurafenib treatment. To this end, we designed multiple RNA FISH probe sets to separately label the cells that required DOT1L inhibition to become resistant and cells that become resistant irrespective of DOT1L inhibitor treatment (Fig. 5a,b). (We expected these probe sets to label fewer than 1:10,000 cells.) We then used these probes to sort corresponding cells from Carbon Copies fixed prior to vemurafenib treatment (Fig. 5c, Supplementary Fig. 12, and Supplementary Fig. 13). RNA sequencing of the sorted subpopulations revealed a few dozen genes differentially expressed between cells that required DOT1L inhibition to survive vemurafenib treatment and non-primed cells (Fig. 5d and Supplementary Fig. 14a-e). Interestingly, we observed differences in expression even in the absence of DOT1L inhibition, suggesting that these genes marked a subpopulation that exists independent of the inhibition of DOT1L, but nevertheless requires DOT1L inhibition in order to become resistant (Supp. Fig. 14). While most differentially expressed genes were also expressed in “conventionally primed” cells, there were a few genes whose expression was somewhat specific to cells that were primed for resistance only when DOT1L was inhibited (Fig. 5d,e and Supplementary Fig. 14a-c). Of these, we selected the gene *DEPTOR*, whose expression we sought to characterize in single cells in our Carbon Copy by RNA FISH (Fig. 5f). (We also chose another gene, *MGP*, whose expression was similarly highly elevated, but only in one replicate; Supplementary Fig. 15.)

**Figure 5:**
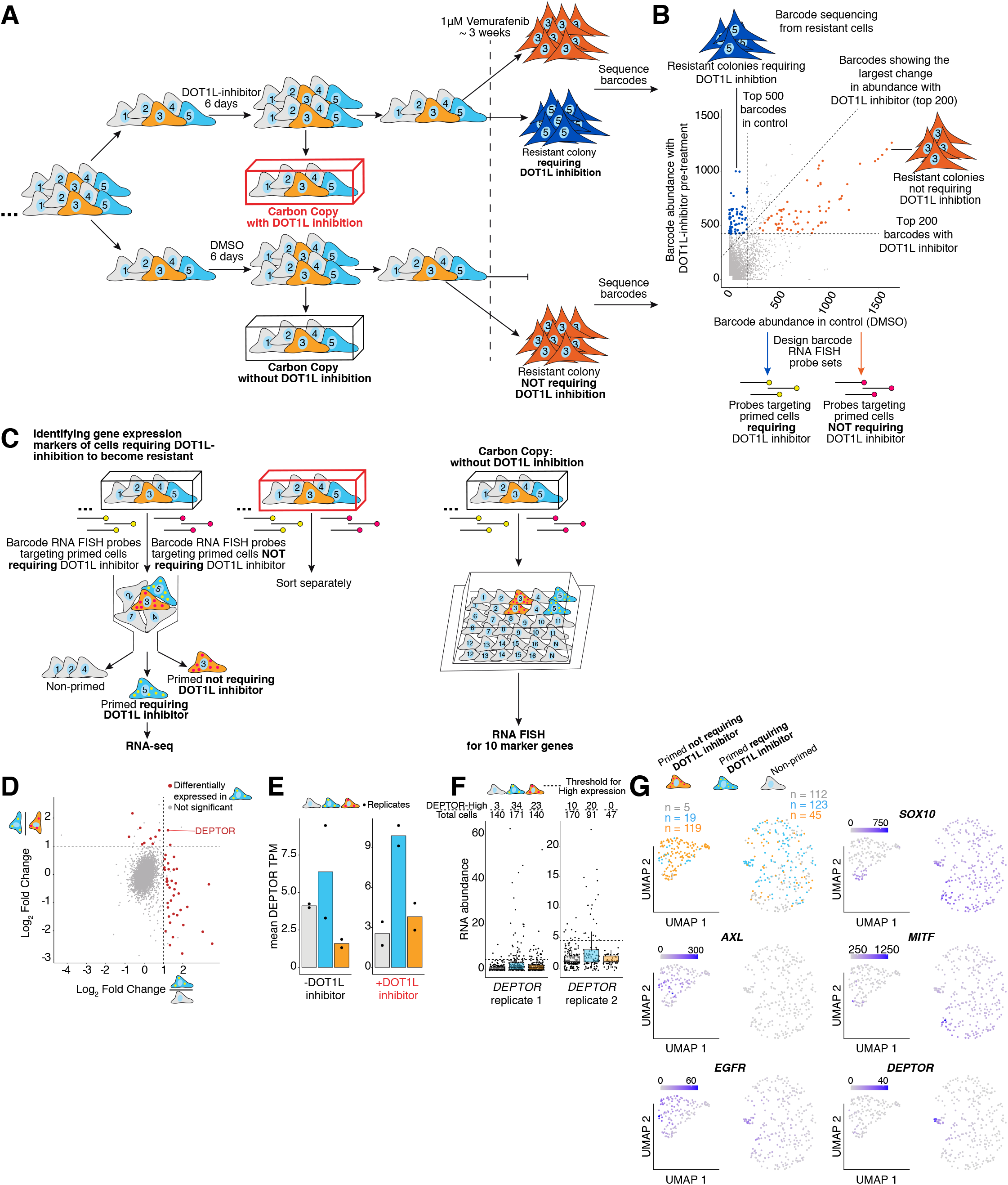
Rewind identifies a distinct subpopulation of cells that require DOT1L inhibition to become vemurafenib resistant. **A**. Experimental approach for identifying the subpopulation of cells that require DOT1L inhibition to become vemurafenib resistant. These experiments began with approximately 400,000 WM989 A6-G3 cells transduced at an MOI ∼ 1.0 and allowed to divide for 10-11 days (∼3-4 population doublings) before splitting the culture into two groups. We treated one group with 4 μM DOT1L inhibitor (pinometostat) and the other with vehicle control (DMSO) for another 6 days (∼2-3 population doublings). We then split each group again, fixing half as our “Carbon Copies” and treating the other half with 1 μM vemurafenib for ∼2.5 weeks. After vemurafenib treatment, we extracted genomic DNA from the remaining cells for barcode sequencing. Note that in principle, DOT1L inhibition may alter cell state (color) even before vemurafenib treatment, which is not depicted here for clarity. **B**. For each barcode identified by sequencing, we plotted its abundance in resistant cells pre-treated with DOT1L inhibitor versus its abundance in resistant cells pre-treated with vehicle control. This comparison revealed a subset of barcodes with a greater relative abundance in resistant cells pre-treated with DOT1L inhibitor (blue points). We used these barcodes to design RNA FISH probes targeting cells that required DOT1L inhibition to become vemurafenib resistant. A separate set of barcodes were highly abundant in resistant cells both with or without DOT1L inhibition (orange points), suggesting that these cells were destined to become resistant whether or not they were pre-treated with DOT1L inhibitor. We used these barcodes to design RNA FISH probes targeting primed cells not requiring DOT1L inhibition to become resistant. Dashed, diagonal line demarcates the 200 barcodes with the largest increase in abundance with DOT1L inhibitor pre-treatment. **C**. Using these probes, we labeled and sorted cells requiring DOT1L inhibition to become vemurafenib resistant (blue), primed cells not requiring DOT1L inhibition (orange), and non-primed cells (gray) from Carbon Copies for RNA sequencing. We separately sorted cells from Carbon Copies treated with DOT1L inhibitor and Carbon Copies treated with vehicle control (2 biological replicates each). **D**. To identify markers of cells that require DOT1L inhibition to become resistant, we used DESeq2 to compare their gene expression to non-primed cells (x-axis) and primed cells not requiring DOT1L inhibition (y-axis). In this analysis, we included cells sorted from all Carbon Copies (treated with DOT1L inhibitor or vehicle control) from 2 biological replicates and included DOT1L inhibitor treatment as a covariate in estimating log_2_ fold changes. Red points correspond to genes differentially expressed in one or both comparisons (p-adjusted ≤0.1 and log_2_ fold change ≥ 1). **E**. Expression of *DEPTOR* in transcripts per million (TPM) in the subpopulations isolated in B. Points indicate TPM values for experimental replicates. **F**. We used the same probe sets as in B. to identify cells in situ in Carbon Copies fixed prior to vemurafenib treatment, then measured single-cell expression of *DEPTOR, MGP, SOX10, MITF*, and 6 priming markers by RNA FISH. Shown is the expression of *DEPTOR* in the indicated cell populations identified in the Carbon Copies treated with vehicle control. Each point corresponds to an individual cell. Above each boxplot is the proportion of cells with levels of *DEPTOR* RNA above the indicated threshold (∼95th percentile in non-primed cells). **G**. We applied the UMAP algorithm to visualize the single-cell expression data from *in situ* Carbon Copies. These plots include 423 cells from the vehicle control treated Carbon Copy. In the upper left plot, points are colored according to the fate of each cell as determined by its barcode, and the number of cells corresponding to each fate are labelled separately above the two largest groupings. For the remaining plots points are colored by the expression level of the indicated gene in that cell. These data correspond to 1 of 2 biological replicates (see Supplementary Fig. 14 for the replicate data).

For single-cell analysis, we performed RNA FISH on the Carbon Copies (half treated with DOT1L inhibitor and half treated with vehicle control) for 10 total genes: 6 priming markers, *SOX10, MITF, DEPTOR*, and *MGP*. We scanned through ∼2 million cells to find those expressing the targeted barcodes, identifying 850 such cells. Using UMAP, we first visualized the expression profiles of cells from the vehicle control treated Carbon Copy, overlaying the information provided by barcode RNA FISH to label cells by their fates (Fig. 5g). We found that the primed cells that did not require DOT1L inhibition to become resistant separated into a distinct grouping that, as before, expressed the previously identified markers such as *AXL* and *EGFR* (Fig. 5g and Supplementary Fig. 14f,g). We initially expected the expression of these genes to also be elevated in the cells that required DOT1L inhibition to become resistant, but perhaps to a lesser extent, reflecting a “subthreshold” state that was unable to survive vemurafenib treatment alone. Contrary to this expectation, the expression profile of this new subpopulation was far more similar to the general population of cells that were not primed for resistance in either condition (Fig. 5g). While in the UMAP projection, many of these cells were grouped together with the non-primed cells, there was a distinct grouping nearby that consisted almost exclusively of cells that were primed for resistance only upon DOT1L inhibition. These cells specifically expressed high levels of *DEPTOR*, along with slightly elevated levels of *EGFR* and lower levels of *MITF*, but showed no differences in the expression levels of the other genes measured compared with non-primed cells (Fig. 5g and Supplementary Fig. 14f-h). (Cells requiring DOT1L inhibition for priming were also enriched for *MGP* in a separate replicate experiment; Supplementary Fig.15.) Taken together, the identification of a unique molecular state marked by *DEPTOR* expression in the overall absence of established priming markers highlights the existence of a qualitatively distinct rare cell state that can lead to drug resistance when a DOT1L inhibitor is given prior to vemurafenib. It is noteworthy that many of the primed cells which require DOT1L inhibition to become vemurafenib resistant expressed neither *DEPTOR* nor established markers (e.g. *AXL, NGFR, ITGA3* etc.) and further work is needed to identify features that better distinguish this rare subpopulation.

While this subpopulation expressed low levels of established priming markers initially, we wondered whether DOT1L inhibition pushed these cells towards a molecular state more similar to the conventional primed cell state (i.e. high levels of *AXL, EGFR, NGFR*, etc.; Fig. 6a). To this end, we compared the transcript levels as measured by RNA sequencing from cells sorted from Carbon Copies treated either with DOT1L inhibitor or vehicle control (Fig. 6b. As expected, with vehicle control, cells that require DOT1L inhibition to become vemurafenib resistant clustered separately from primed cells that do not require DOT1L inhibition (Fig. 6c,d). With DOT1L inhibition, these two populations appeared modestly more similar transcriptionally, however they remained predominantly distinct (Fig. 6c,d). RNA FISH on cells that require DOT1L inhibition to become resistant revealed that DOT1L inhibition did not increase expression of established priming markers, and if anything, modestly decreased their expression (Fig. 6e,f and Supplementary Fig. 16a,b). Overall, these gene expression differences between primed subpopulations both before and after DOT1L inhibition suggest that DOT1L inhibition does not simply convert cells into the previously established primed cell state capable of surviving vemurafenib treatment, but rather, it may reveal a separate route to resistance.

**Figure 6:**
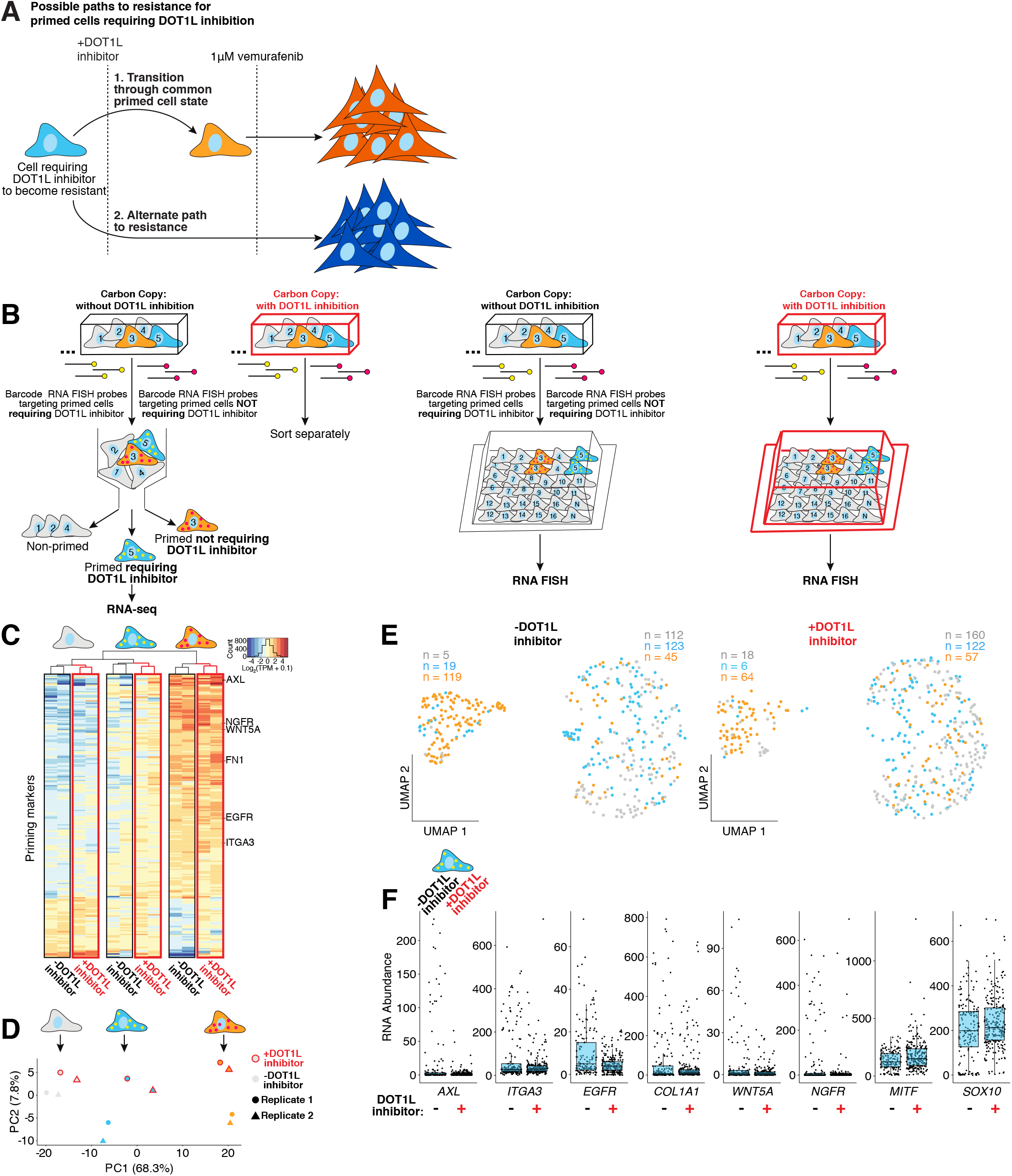
DOT1Li inhibition enables a new subpopulation of cells to survive vemurafenib treatment without converting them into the known primed cell state. **A**. We asked whether DOT1L inhibition enables a new subpopulation of cells to survive vemurafenib treatment by converting them into the previously established primed cell state or whether these cells become resistant via a possible alternative path. **B**. We used Rewind to isolate and perform RNA sequencing on cells requiring DOT1L inhibition to survive vemurafenib treatment (blue), cells not requiring DOT1L inhibition (orange), and non-primed cells (gray) sorted from both Carbon Copies treated with DOT1L inhibitor (red outline) and Carbon Copies treated with vehicle control (gray outline) (2 replicates each sorted for RNA sequencing). **C**. Heatmap displays expression of established priming markers across sorted subpopulations from control and DOT1L-inhibitor pre-treated Carbon Copies. Dendrogram shows hierarchical clustering of samples by expression values. We defined priming markers as protein-coding genes differentially expressed (p-adjusted ≤ 0.1 and abs(log_2_ fold change) ≥ 1) in primed cells not requiring DOT1L inhibition versus non-primed cells isolated from the Carbon Copy treated with vehicle control. **D**. Using expression of priming markers as in C., we performed principal component analysis on primed and non-primed cell populations. Red outline indicates samples sorted from the Carbon Copy treated with DOT1L inhibitor. **E**. We used the same probes as in B. to identify cell populations in situ in Carbon Copies treated with DOT1L inhibitor or vehicle control. We then used RNA FISH to measure single-cell expression of several established priming markers and visualized the relationship in gene expression between single cells using the UMAP algorithm with the first 6 principal components. This analysis included expression data from 850 single cells. Points are colored according to the fate of each cell as determined by its barcode, and the number of cells corresponding to each fate are labelled above the largest groupings. **F**. Plotted are single-cell expression data for 6 priming markers, *MITF* and *SOX10* in cells that require DOT1L inhibition to become vemurafenib resistant. Each point corresponds to an individual cell. Below each boxplot, we indicate whether the cells are from the Carbon Copy treated with DOT1L inhibitor (+) or vehicle control (-). The corresponding data for non-primed cells and primed cells not requiring DOT1L inhibition are shown in Supplementary Fig. 16. These data correspond to 1 biological replicate.

## Discussion

We have here revealed the existence of a rich set of rare subpopulations within seemingly homogenous cells, several of which can lead to phenotypically distinct fates. Despite the population having a clonal origin and being grown in homogeneous cell culture conditions, these subpopulations spontaneously emerge via transient cell-state fluctuations that can persist for several cell divisions. It remains unclear how precisely these subpopulations arise, although, intriguingly, it may arise from network interactions between multiple regulatory factors ^28^. It is also unclear how these states revert to the population baseline. We here observe states persisting for over 5-6 generations, whereas previous reports based on sorting by individual markers suggested reversion on shorter timescales ^6^. It is possible that the more pure primed population identified by Rewind can persist longer than impure populations which may contain transient intermediates.

For the variability that is associated with priming, it is tempting to imagine single axes of variability for both state and fate, in which cells that have fluctuated further up a putative primed state hierarchy lead to different degrees of resistance. However, our results show that even for the simple case of heterogeneity in the size of resistant clones, expression of the rare cell markers *AXL/ITGA3/EGFR* and *WNT5A*/*MMP1/NGFR* varied along at least two axes prior to the addition of drug, with each axis being associated with either the low-abundance or high-abundance clones. Further use of tools like Rewind, potentially in combination with transcriptome-scale RNA FISH or single-cell RNA sequencing, may help to fully reveal the structure of these fluctuations and consequent subpopulations. Resistant cell fates likely have similarly complex modes of variability, and our results suggest that these modes likely have origins in molecular variability in the initial cell state. The nature of these mappings may help guide therapy, and it may be important to consider the multiple different initial primed cellular states that give rise to resistant cells following distinct treatments, as highlighted by our DOT1L inhibition results.

A critical consideration in developing Rewind was minimizing contamination from “off-target” non-primed cells. These cells could in principle come from probes falsely generating signal in non-primed cells or technical limitations of FACS. These contaminating cells can dramatically dilute measurements of gene expression specific to the targeted, rare subpopulations (Supplementary Fig. 1d,e). We found that barcode detection by FACS was far more prone to contamination than barcode detection by imaging, which had very high precision (estimated to be ∼97%; Supplementary Fig. 1f-h); indeed, we believe it is for this reason that we observe larger magnitude differences by RNA FISH than by RNA sequencing of sorted populations, particularly for markers down-regulated in primed cells such as *SOX10* and *MITF*. Yet, despite these concerns, we discovered and validated the priming markers *ITGA3* and *AXL*, while also identifying previously known markers such as *NGFR* and *EGFR*. We also found that experiment-to-experiment technical variability was relatively minimal: by imaging, we did not see much difference in off-target signal across different probe sets (with rare exceptions of “dirty” probes), and barcode sequencing of cDNA from sorted subpopulations labelled with different probe sets suggested similar levels of enrichment (Fig. 1, Supplementary Fig. 12 and a notable exception in Supplementary Fig. 6)).

The global transcriptional profiles afforded by RNA sequencing of rare primed cells allowed us to ask what pathways might be active in these cells beyond the ones like growth factor receptor signaling that have already been associated with vemurafenib resistance in melanoma ^6,25,29-31^. One of the strongest signatures was the upregulation of cell adhesion proteins and structural components of the extracellular matrix. Such signatures suggest the possibility that control of cell state and behavior may have both a component that is autonomous to the cell itself and a component that is instructed by the extracellular matrix. Future research may help reveal if and how the extracellular matrix is able to influence primed cellular states, and consequently, therapy resistance.

There were also several other expression signatures active in distinct subpopulations of cells. For instance, *DEPTOR* expression marked one set of primed cells. While *DEPTOR* may not have any functional role in priming, it is known that DEPTOR inhibits mTOR signaling, which may relieve negative feedback on PI3K/Akt signaling, and, in turn, bypass the inhibition of BRAF signaling ^32^. Further work is needed to establish such potential mechanisms.

The processes involved in the acquisition of stable drug resistance act both on short timescales (such as signaling) and on longer timescales (transcription). For instance, vemurafenib acts by inhibiting MAPK signaling, but the vemurafenib treatment itself relieves negative feedback on growth factor receptor signaling and allows ERK reactivation via BRAF^V600E^-independent routes ^27,33^. Single-cell analysis of ERK signaling has shown that individual cells vary dramatically in ERK activity following vemurafenib treatment with rare cells reactivating ERK to levels comparable to untreated cells. Rewind allowed us to connect these near-term single-cell signaling dynamics in rare cells to both their initial transcriptional state and their ultimate resistant fate. These connections revealed that the primed melanoma cells that go on to survive vemurafenib treatment had both higher levels of phosphorylated ERK soon after treatment and expressed multiple receptor tyrosine kinases along with their cognate ligands. It is possible that this unique gene expression program enabled autonomous ERK reactivation.

We chose to focus on the priming of melanoma cells towards different fates following targeted therapy treatment. However, there are several examples in which non-genetic differences can lead rare cells to undergo important transformations, including the induction of pluripotency in otherwise terminally differentiated cells ^34^ and transdifferentiation of one cell type into another. Application of techniques like Rewind in these contexts may reveal universal characteristics of priming and reprogramming.

## Acknowledgements

We thank C. Bartman, A. Anguierra, J. Murray, N. Zhang, L. Cai, B. Stanger, A. Coté, K. Kiani, E. Sanford, and Y. Goyal along with other members of the Raj lab for many useful suggestions. We thank the Flow Cytometry Core Laboratory at the Children’s Hospital of Philadelphia Research Institute for assistance in designing and performing FACS, including F. Tuluc for several helpful discussions. BLE acknowledges support from NIH training grants F30 CA236129, T32 GM007170 and T32 HG000046, EAT acknowledges R01 CA238237, IPD acknowledges NIH 4DN U01 HL129998, NIH Center for Photogenomics (RM1 HG007743), CLJ acknowledges NIH T32 DK007780 and F30 HG010822, NJ acknowledges NIH F30 HD103378, SMS acknowledges support from DP5 OD028144 AR acknowledges R01 CA238237, NIH Director’s Transformative Research Award R01 GM137425, R01 CA232256, NSF CAREER 1350601, P30 CA016520, SPORE P50 CA174523, NIH U01 CA227550, NIH 4DN U01 HL129998, NIH Center for Photogenomics (RM1 HG007743), and the Tara Miller Foundation.

## Author Contributions

BLE and AR conceived of and designed the project with input from EAT and SMS. BLE performed experiments and analyses. IPD, CLJ and NJ assisted with optimizing barcode RNA FISH protocols. CC assisted with cell and colony segmentation. AR supervised the project. BLE and AR wrote the manuscript with helpful feedback from SMS, CLJ, NJ and other members of the Raj lab.

## Materials and Methods

### Barcode Lentivirus Library Construction

Starting with the LRG2.1T plasmid kindly provided by Dr. Junwei Shi, we derived a lentivirus vector backbone for Rewind by removing the U6 promoter and sgRNA scaffold then inserting a spacer sequence flanked by EcoRV restriction sites after the stop codon of GFP. For the barcode insert, we ordered PAGE-purified Ultramer oligonucleotides (IDT) containing “WSN” repeated for 100 nucleotides (W=A or T, S = G or C, N = Any) flanked by 30 nucleotides homologous to the vector insertion site for Gibson Assembly (see Supplementary Table 1 for barcode insert sequence). We then digested the vector backbone overnight with EcoRV (NEB), gel purified the linearized vector. We combined 100ng of linearized vector, 1.08 μL barcode oligo insert (100 nM in nuclease-free water), 10 μL Gibson assembly master mix (NEB E2611) and nuclease free water to a final volume of 20 μL then incubated the reaction at 50°C for 1 hour. We next column purified the assembled plasmid using Monarch DNA cleanup columns (NEB) according to the manufacturer’s protocol then electroporated 2 μL of the column purified plasmid into Endura Electrocompetent *E. coli* cells (Lucigen) using a GenePulserXCell (Biorad) with the following settings: 25msec pulse length, 10 μF capacitance, 600Ω resistance, and 1800V voltage. We performed 6 electroporations using the same plasmid in parallel. Immediately after electroporation, we added 1 mL of pre-warmed (37°C) recovery media to each electroporation cuvette then transferred the liquid to 1.5 mL microcentrifuge tubes and placed these tubes on a shaker at 225rpm and 37°C for 1 hour. After this recovery, we took 10 μL of the culture for plating serial dilutions and transferred the rest to 150-200 mL of 1X LB Broth containing 100 μg/mL ampicillin. We incubated these cultures on a shaker at 225 rpm and 32°C for 12-14 hours then pelleted the cultures by centrifugation and isolated plasmid using the EndoFree plasmid maxiprep kit (Qiagen) according to the manufacturer’s protocol. In some instances, pellets were frozen at −20°C for several days before plasmid isolation. To estimate transformation efficiency, we counted colonies on the plated serial dilutions and verified barcode insertion by PCR from 20-30 colonies per plate. We pooled the plasmids from the 6 separate cultures in equal amounts by weight before packaging into lentivirus. This protocol is also available online at https://www.protocols.io/view/barcode-plasmid-library-cloning-4hggt3w

### Cell Lines and Culture

We derived the WM989 A6-G3 melanoma cell line by twice single-cell bottlenecking the WM989 melanoma cell line kindly provided by Dr. Meenhard Herlyn (Wistar Institute) ^6,35^. Similarly, we derived WM983b E9-C6 by twice single-cell bottlenecking the WM983b melanoma cell line also provided by Dr. Meenhard Herlyn. We verified the identity of these cell lines by DNA STR Microsatellite fingerprinting at the Wistar Institute.

We cultured both melanoma cell lines in TU2% media consisting of 80% MCDB 153, 10% Leibovitz’s L-15, 2% FBS, 2.4 mM CaCl_2_, 50 U/mL penicillin, and 50 μg/mL streptomycin and passaged cells using 0.05% trypsin-EDTA. For harvesting drug-treated resistant cells we used 0.1% Trypsin-EDTA. For lentivirus packaging, we cultured HEK293FT cells in DMEM containing 10% FBS 50 U/mL penicillin and 50 μg/mL streptomycin and passaged cells using 0.05% Trypsin-EDTA.

### Lentivirus Packaging and Transduction

Prior to plasmid transfection, HEK293FT cells were grown to ∼90% confluency in 6-well plates in DMEM containing 10% FBS without antibiotics. For each 6-well plate, we added 80 μL PEI to 0.5 mL Opti-MEM (ThermoFisher 31985062) and separately, combined 7.5 μg pPAX2, with 5 μg VSVG and 7.71 μg of the barcode plasmid library in 0.5 mL Opti-MEM then incubated the solutions at room temperature for 5 minutes. We then mixed the 2 solutions together with vortexing and incubated the combined solution at room temperature for 15 minutes. We added 184 μL of the plasmid-PEI solution dropwise to each well of the 6-well plate. After 6-8 hours, we aspirated the media from the cells, washed the cells once with 1X DPBS, then added fresh culture media (DMEM containing 10% FBS and antibiotics). The following morning, after confirming that the majority of cells were GFP positive, we aspirated the media, washed the cells once with 1X DPBS then added 1 mL of TU2% to each well. Approximately 12 hours later, we transferred the virus laden media to a falcon tube and added another 1 mL of TU2% to each well. We collected virus laden media twice more over the next ∼16 hours and during this time, stored the collected media at 4°C. After the final collection, we filtered the virus laden media through a 0.22 μm PES filter then stored 1-2 mL aliquots at −80°C.

To transduce WM989 A6-G3 and WM983b E9-C6 cells we added freshly thawed (on ice) virus laden media and polybrene (final concentration 4μg/mL) to dissociated cells, then plated the cells onto 6-well plates (100,000 cells in 2 mL media per well) and centrifuged the plate at 1,750 rpm (517 × g) for 25 minutes. We incubated the cells with virus for 6-8 hours then removed the media, washed the cells once with 1X DPBS and added 3mL of TU2% to each well. The following day, we passaged the cells to 10 cm dishes (one 6-well plate into three 10 cm dishes). For WM989 A6-G3, we split barcoded cells into Carbon Copy and separate vemurafenib treatment groups 11 days after transduction for sort experiments (Fig. 1) or 10 days after transduction for in situ experiments (Fig. 2-4) unless otherwise specified. These timepoints correspond to 4-5 population doublings since transduction. For WM983b E9-C6, we split barcoded cells into Carbon Copy and separate vemurafenib treatment groups 7 days after transduction (also corresponding to 4-5 population doublings) unless otherwise specified. We cultured in situ Carbon Copies for 4 days before fixation in order to more easily identify clusters of cells expressing targeted barcodes.

### Simulations of Experimental Conditions Used for Rewind

As described above, we expanded barcoded cells for at least 4 population doublings before splitting-off the Carbon Copy and drug-treatment groups for Rewind. As such, there were on average ∼16 closely-related cells for each barcoded clone before the split. For a 50:50 split, the probability that at least 1 of 16 cells ends up in both groups is ∼99.997%, or in other words, less than 0.002 % of clones are expected to be “missing” from either group. However, given the unavoidable variability in cell growth, it is likely that some clones will have divided fewer than 4 times, and these clones are more likely to be entirely missing from the Carbon Copy. (We note that we do not care about clones that are missing from the drug treatment group since they will not become resistant colonies and their barcodes will not be selected for probe design). To empirically estimate the proportion of clones present in our Carbon Copy, we sequenced barcode gDNA from barcoded WM989 A6-G3 after ∼4 population doublings, then computationally “split” the sequenced barcodes into 2 halves, after first weighting each barcode by its read count and scaling the average read count to 16. Finally, we calculated the proportion of barcodes present in both halves. Simulating this procedure 10,000 times, we found that ∼92.3-92.6% of barcodes were present in both halves and <4% of barcodes were “missing” from the simulated Carbon Copy.

We also note that to eliminate spurious barcodes arising due to PCR or sequencing errors, we merged highly similar barcode sequences as described further below (see **Computational analyses of barcode sequencing data**) and filtered barcodes with fewer than 5 unique reads. The simulations were robust to a range of read count thresholds ≥ 2.

We used the same barcode sequencing data to simulate the “heritability-split-experiment” for Supplementary Fig. 3d. In this case, we randomly sampled 200 barcodes twice (without replacement and weighting each barcode by its read count), then calculated the proportion of barcodes shared between the two samples. We performed the same simulation for WM983b E9-C6 (Supplementary Fig. 6b) using sequencing data from barcoded WM983b E9-C6 grown for ∼4 population doublings.

The scripts used for these simulations are available on Dropbox at https://www.dropbox.com/s/p5t9onmezasmtty/heritabilitySplitWM989.R?dl=0.

### Fluorescence Activated Cell Sorting (FACS)

To isolate ITGA3-High WM989 A6-G3, we first trypsinized and pelleted 8 confluent 10cm plates, washed once with 1X DPBS containing 0.1% BSA (0.1% BSA-PBS), and then split the cells into two equal pellets. We resuspended each pellet in 0.4 mL 0.1% BSA-PBS containing 1:200 anti-ITGA3 antibody (DSHB clone P1B5 stock concentration 354 μg/mL) then incubated on ice for 1 hour. After primary incubation, we pelleted the cells, washed twice with ∼5 mL 0.1% BSA-PBS then resuspended cells in 0.16 mL 0.1% BSA-PBS containing 1:500 anti-mouse FAb2 secondary antibody conjugated to Alexa Fluor 488 (Cell Signaling #4408) and incubated on ice for 30 minutes. Finally, we pelleted the cells, washed twice with 0.1% BSA-PBS, then resuspended the pellet in 0.1% BSA-PBS containing 100 ng/mL DAPI and proceeded with FACS on a MoFlo Astrios (Beckman Coulter). After gating for singlets and live cells, we collected 15,000 events from the brightest 0.3-0.4% ITGA3-High gate and equal numbers from the dimmest ∼99% ITGA3-Low gate. We plated two thirds of the sorted cells onto 2-well glass bottom chamber plate (Nunc Lab-Tek 155380) for treating with vemurafenib (see below) and the rest on a separate 2-well glass bottom chamber plate for verifying *ITGA3* expression by single-molecule RNA FISH.

We followed a similar procedure for isolating AXL-High WM983b E9-C6 starting with 10 10cm dishes split into two equal cell pellets, performing all incubations and washes with 1% BSA-PBS and staining with 1:50 primary antibody (goat anti-human AXL AF154 from Novus Biologicals) and 1:60 secondary antibody (bovine anti-goat conjugated to Alexa Fluor 647; Jackson ImmunoResearch 805-605-180). After gating for singlets and live cells, we collected 20,000 events from the brightest ∼0.3% AXL-High gate and equal numbers from the dimmest ∼20% AXL-Low gate, then plated cells onto 2-well glass bottom plates (10,000 cells per well) for vemurafenib treatment or RNA FISH as above.

### Drug Treatment Experiments

We prepared stock solutions of 4mM vemurafenib (PLX4032, Selleckchem, S1267), 10mM pinometostat (SelleckChem S7062), 100 μM trametinib (SelleckChem S2673), and 10mM Dabrafenib (SelleckChem S2807). We prepared all stock solutions in DMSO and divided into small aliquots stored at −20°C to minimize freeze-thaw cycles. For drug treatment experiments, we diluted the stock solutions in culture medium to a final concentration of 1 µM for vemurafenib, 4 µM for pinometostat, 10 nM for trametinib, and 1 µM for dabrafenib unless otherwise specified.

For Rewind experiments in WM989 A6-G3, we treated cells for 3 weeks replacing media containing drug every 3-4 days. Following vemurafenib treatment, we trypsinized and collected all remaining cells, washed cells once with 1x DPBS, then pelleted and froze 90% of the cells at −20°C until gDNA extraction and barcode sequencing as described below. We fixed the remaining 10% of vemurafenib resistant cells for barcode RNA FISH, FACS and RNA sequencing as described below and in Supplementary Fig. 5a. For DOT1L inhibitor pre-treatment, we treated cells with 4 µM pinometostat for 6 days, replacing media on day 3 and again when splitting off the Carbon Copy on day 5. Following the ITGA3 sort, we fixed WM989 A6-G3 cells after 18 days of vemurafenib treatment in order to more easily quantify numbers of colonies. For Rewind experiments in WM983b E9-C6, we treated cells for 4 weeks replacing media containing 1 µM of vemurafenib every 3-4 days. Cells surviving drug treatment were harvested and frozen as described above.

### Cell Quantification

Following drug treatment experiments, we fixed cells by incubation for 10 minutes in 3.7% formaldehyde (Sigma F1635) diluted in 1X PBS, followed by two washes with 1X PBS then overnight permeabilization at 4°C with 70% ethanol. We stained nuclei by incubation in 2X SSC containing 50 ng/mL DAPI then imaged the majority of each well via a tiling scan at 20X magnification. To quantify cell and colony numbers, we used custom MATLAB software to stitch the tiled images, identify nuclei and manually circle individual resistant colonies. Software and scripts used for these analyses can be found: https://github.com/arjunrajlaboratory/colonycounting_v2 and https://www.dropbox.com/sh/p279h7mak0rrklx/AACyM_IiVP3prkjdDmd6HqOca?dl=0.

### Barcode Library Preparation and Sequencing

We isolated genomic DNA (gDNA) from barcoded cells using the QIAmp DNA Mini Kit (Qiagen 51304) according to the manufacturer’s protocol. We performed targeted amplification of the integrated barcode vector using custom primers containing Illumina adapter sequences, unique sample indexes, variable length staggered bases, and 6 random nucleotides (“UMI”; NHNNNN) which, despite not uniquely tagging barcode DNA molecules, appeared to modestly increase reproducibility between replicate libraries and normalize read counts (see Supplementary Table 2 for a complete list of primers). For each sample, we performed multiple PCR reactions (using 20-40% of the total isolated gDNA) each consisting of 1 μg of gDNA, 500 nM primers, 25 μL NEBNext Q5 HotStart HiFi PCR master mix and nuclease free water to a final volume of 50 μL. We ran the reactions on a thermal cycler with the following settings: 98°C for 30 seconds, followed by N cycles of 98°C for 10 seconds then 65°C for 40 seconds, and finally 65°C for 5 minutes. After the PCR, we purified libraries using 35 μL (0.7X) Ampure XP magnetic beads with two 80% ethanol washes followed by final elution in 20 μL 0.1X TE (1 mM Tris HCl pH 8.0 100 μM EDTA). Purified libraries from the same sample were pooled, quantified using the Qubit dsDNA High Sensitivity assay (ThermoFisher) then sequenced on a NextSeq 500 using 150 cycles for read 1 and 8 cycles for each index. For barcoding experiments not requiring RNA FISH probe design, shorter reads (75 cycles) provided sufficient information to identify unique barcodes.

To reduce PCR amplification bias, we determined the number of cycles (“N”) for each sample by first performing a separate qPCR reaction and selecting the number of cycles needed to achieve ⅓ of the maximum fluorescence intensity. We included 0.25 μL 100X SYBR Green I (10,000X diluted 1:100 in 10 mM Tris pH 8.0; Invitrogen S7563) per 25 μL qPCR reaction and, when possible, performed multiple reactions with serial dilutions of gDNA (1:4 and 1:16). For experiments with multiple similar samples (same MOI, same treatment) we performed qPCR on one of these samples and extrapolated “N” to the rest.

To test reproducibility of our barcode quantification, for a subset of samples we prepared duplicate libraries with separate indexes and compared barcode read counts between these technical replicates. As shown in Supplementary Fig. 2, we found a high correlation (>95%) in barcode abundance between these technical replicates.

### Computational Analyses of Barcode Sequencing Data

We recovered barcodes from sequencing data using custom Python scripts available at: https://github.com/arjunrajlaboratory/timemachine. These scripts search through each read to identify sequences complementary to our library preparation primers, and if these sequences pass a minimum length and phred score cutoff, then the intervening barcode sequence is counted. In addition to counting total reads for each barcode, we also count the number of “UMIs” incorporated into the library preparation primers (see above). While we do not believe that these “UMIs” tag unique barcode DNA molecules, empirically they appeared to slightly improve the correlation in barcode abundance between replicate libraries and were therefore used for most subsequent analyses. Using the STARCODE software ^36^ (available at https://github.com/gui11aume/starcode), we merged highly similar barcode sequences (Levenshtein distance ≤ 8), summing the counts and keeping only the more abundant barcode sequence.

For selecting barcodes corresponding to resistant colonies, we ranked the barcode sequences by counts then converted the most abundant 100-200 barcodes sequences into fasta files for probe design as described below. Barcode sequences with ≥30 bases of homology to the vector backbone were excluded for concerns of generating non-specific FISH probes (we checked for non-specific binding a second time during probe design as described below).

We selected barcodes corresponding to resistant colonies that require DOT1L inhibition using the following criteria: 1. Among the most abundant 200 barcodes in DOT1L inhibitor pre-treated resistant cells, 2. not among the most abundant 500 barcodes in the DMSO pre-treated resistant cells and 3. greatest difference in abundance between DOT1L inhibitor pre-treated and DMSO pre-treated resistant cells among all barcodes passing criteria 1 and 2. For barcodes corresponding to resistant colonies not requiring DOT1L inhibition, we selected sequences that were: 1. in the top 200 barcodes in both the DOT1L-inhibitor and DMSO pre-treated resistant cells and 2. which had relatively similar abundances across these two conditions (not among the 500 barcodes with the largest difference in abundance).

### Barcode RNA FISH Probe Design

Using fasta files of selected barcodes, we design HCR probes using Rajlab ProbeDesignHD software(code freely available for non-commercial use here https://flintbox.com/public/project/50547/). For each barcode sequence, we designed 2 non-overlapping 42mer probes with a target Gibbs free energy for binding of −55 (allowable Gibbs Free Energy [-65, −45]). We excluded probes with complementarity to repetitive elements, pseudogenes or the vector backbone used to generate the barcode plasmid library. We then split each 42mer probe into 2 20mer sequences (removing the middle two nucleotides) and appended split-initiator HCR sequences using custom python scripts (see Supplementary Table 3 for sequences) ^37^. For each 20mer sequence, we measured the maximum complementarity to the vector backbone and other barcodes present in the sample in order to manually exclude probes with potential for non-specific hybridization. We ordered the final probe sequences synthesized from IDT in picomole scale 384 well plates. Finally, we resuspended barcode HCR probes to 50 μM in nuclease-free water then combined these probes into pools each containing 24 different barcode probes at a final concentration of 2 μM each.

For ClampFISH we designed 30mer probes targeting select barcodes using Rajlab ProbeDesignHD software with a target Gibbs free energy of −40 (allowable Gibbs Free Energy [-50, −30]). As above, we excluded probes with complementarity to repetitive elements, pseudogenes or the vector backbone. We then appended 10mer sequences to the 5’ and 3’ ends of each probe (used for subsequent ligation) and ordered the final probe sequences synthesized from IDT in picomole scale 384 well plates. We resuspended barcode ClampFISH probes to 100 μM in nuclease-free water then combined these probes into pools each containing 30 different barcode probes. To these pools we ligated oligonucleotides (oligos) containing alkyne and azide modifications at their 5’ and 3’ ends, respectively (see Supplementary Table 4 for sequences). For this ligation, we first phosphorylated the 5’ ends of each probe set by combining 4 μL of the pooled oligos with 1 μL T4 PNK (NEB), 20 μL T7 DNA ligase reaction buffer (NEB), and 2 μL nuclease-free water then incubating at 37°C overnight. Next, we added the alkyne and azide modified oligos along with complementary bridging 20mer oligos (3 μL each of 400 μM stocks) and heated the reactions to 95°C for 5 minutes then cooled to 12° C at a rate of −0.1° C/second. After cooling, we added 1 μL T7 ligase (NEB) and incubated overnight at room temperature. We purified the ligated barcode ClampFISH probes using Monarch DNA cleanup columns (NEB) according to the manufacturer’s protocol. This protocol for generating barcode clampFISH probes is also available online at https://www.protocols.io/view/invertedclampfish-ligation-qxwdxpe. We prepared amplifier probes MM2B, MM2C, P9B and P9C as described previously ^38^.

### RNA FISH

We designed oligonucleotide probe sets complementary to our genes of interest using custom probe design software written in MATLAB and ordered them with a primary amine group on the 3’ end from Biosearch technologies (see Supplementary Table 5 for probe sequences). For each gene, we pooled their complementary oligos and coupled the probe set to either Cy3 (GE Healthcare), Alexa Fluor 594 (Life Technologies), or Atto647N (Atto-Tec)NHS ester dyes. We performed single-molecule RNA FISH as described in ^39^ and ^6^ for multiple cycles of hybridization. We aspirated media from adherent cells, washed the cells once with 1X PBS, then incubated the cells in fixation buffer (3.7% formaldehyde in 1X PBS) for 10 minutes at room temperature. We next aspirated the fixation buffer, washed samples twice with 1X PBS, then added 70% ethanol and stored samples at 4° C. For hybridization, we first washed samples with washing buffer (10% formamide in 2X SSC) then applied the RNA FISH probes in hybridization buffer (10% formamide and 10% dextran sulfate in 2X SSC). We covered samples with coverslips then hybridized samples overnight in humidified containers at 37°C. The following morning, we washed samples 2 × 30 minutes with washing buffer at 37°C, adding 50 ng/mL DAPI to the second wash to stain the nuclei. After these washes, we rinsed samples once with 2X SSC then added new 2X SSC and proceeded with imaging. To strip RNA FISH probes, we incubated samples in stripping buffer (60% formamide in 2X SSC) for 20 minutes on a hot plate at 37°C, washed samples 3 × 15 minutes with 1X PBS on a hot plate at 37°C, then returned samples to 2X SSC. After stripping RNA FISH probes, we re-imaged all previous positions and excluded dyes with residual signal from subsequent hybridization.

### Barcode RNA HCR

We adapted the Hybridization Chain Reaction (HCR V3.0)^37^ for barcode RNA FISH as follows. We used 1.2 pmol each of up to 240 barcode RNA FISH probes per 0.3 mL hybridization buffer. Our primary hybridization buffer consisted of 30% formamide, 10% dextran sulfate, 9 mM citric acid pH 6.0, 50 μg/mL heparin, 1X Denhardt’s solution (Life Technologies 750018) and 0.1% tween-20 in 5X SSC. For primary hybridization, we used 100 μL hybridization buffer per well of a 6 well plate, covered the well with a glass coverslip, then incubated the samples in humidified containers at 37°C for 6 hours. Following the primary probe hybridization, we washed samples 4 × 5 minutes at 37°C with washing buffer containing 30% formamide, 9 mM citric acid pH 6.0, 50 μg/mL heparin, and 0.1% tween-20 in 5X SSC. We then washed the samples at room temperature 2 × 5 minutes with 5X SSCT (5X SSC + 0.1% Tween-20), then incubated the samples at room temperature for 30 minutes in amplification buffer containing 10% dextran sulfate and 0.1% Tween-20 in 5X SSC. During this incubation, we snap-cooled individual HCR hairpins (Molecular Instruments) conjugated to either Alexa Fluor 647 (Alexa647), Alexa Fluor 594 (Alexa594) or Alexa Fluor 546 (Alexa546) by heating to 95°C for 90 second then immediately transferring to room temperature to cool for 30 minutes concealed from light. After these 30 minutes, we resuspended and pooled the hairpin in amplification buffer to a final concentration of 6nM each. We added the hairpin solution to samples along with a coverslip, then incubated samples at room temperature overnight (12-16 hours) concealed from light. The following morning, we washed samples 5 × 5 minutes with 5X SSCT containing 50 ng/mL DAPI, added SlowFade antifade solution (Life Technologies S36940) and a coverslip then proceeded with imaging. To remove fluorescent signal for subsequent rounds of RNA FISH or immunofluorescence, we photobleached samples on the microscope or stripped HCR hairpins as described above for RNA FISH probes. We used this modified HCR V3.0 protocol for labeling barcode RNA in all experiments except those indicated in Supplementary Fig. 8, which relied on the ClampFISH protocol described below.

For performing HCR in suspension, we adapted the published protocol ^37^ as follows. We fixed dissociated cells in suspension by washing the cells with 1X DPBS, resuspending the cell in ice cold 1X DPBS, adding equal volume of ice-cold fixation buffer (3.7% formaldehyde 1X PBS) then incubating with rotation at room temperature for 10 minutes. We next pelleted fixed cells by centrifugation at 800 × g for 3 minutes, washed twice with ice cold 1X PBS, then resuspended in 70% ethanol and stored fixed cells at 4°C. For primary probe hybridization we used 0.5 mL hybridization buffer containing 4 nM of each barcode RNA FISH probe and incubated samples using the same conditions as described above. After primary probe hybridization, we washed samples 4 × 10 minutes with 0.5 mL washing buffer then 2 × 10 minutes with 0.5 mL 5X SSCT. We next incubated samples for 30 minutes in amplification buffer and snap-cooled HCR hairpins as described above. For amplification, we used 15 nM final concentration of each HCR hairpin and incubated samples at room temperature overnight concealed from light. After amplification, we washed samples 6 times with 5X SSCT the proceeded with FACS. In between hybridizations and washes, we pelleted cells by centrifugation at 400 × g for 5 minutes and used low-molecular weight dextran sulfate (Sigma D4911) in hybridization and amplification buffers to improve pelleting.

We note that the final hairpin concentrations used in these experiments is 4- to 10-fold lower than the manufacturer’s protocol, which we optimized to reduce nonspecific amplification while still enabling sensitive barcode RNA detection at 20X magnification. At the same time we have noticed lot to lot variation in HCR hairpins purchased from Molecular Instruments with each lot requiring some testing and optimization for use with Rewind. Finally, we found that hybridization and wash buffers without citric acid, heparin, Denhardt’s solution or tween-20 (that is using only SSC, formamide and dextran sulfate) appeared to work as well as the manufacturer’s recommended buffers for barcode RNA HCR and we used these minimal buffers for barcode detection prior to immunofluorescence (Fig. 3).

### Barcode RNA ClampFISH

For Supplementary Fig. 8, we adapted the published ClampFISH protocol ^38^ for labeling barcode RNA as follows. We generated modified primary probes and amplifier probes as described in **Barcode RNA FISH Probe Design**. For hybridization, we washed fixed samples with washing buffer containing 40% formamide in 2X SSC then applied the primary ClampFISH probes in primary hybridization buffer containing 40% formamide, 10% dextran sulfate, 1 mg/mL yeast tRNA (Invitrogen 15401029), 0.02% BSA, and 100 μg/mL sonicated salmon sperm DNA (Agilent 201190-81) in 2X SSC. We included up to 180 ClampFISH probes targeting up to 60 different barcode RNA sequences per hybridization (total probe concentration 125 ng/µL − 250 ng/μL). We added coverslips to samples then hybridized for 6-8 hours in humidified containers at 37°C. After hybridization, we added wash buffer containing 40% formamide in 2X SSC to dislodge coverslips then replaced the wash buffer and incubated the samples for 20 minutes at 37°C. We performed a second wash for 20 minutes at 37°C using buffer containing 20% formamide and 2X SSC then performed the second round of hybridization with MM2B and MM2C amplifier probes in amplifier hybridization buffer (20% formamide, 10% dextran sulfate, 1 mg/mL yeast tRNA, 0.02% BSA, and in 2X SSC.; final probe concentration 10 ng/μL each). After the second hybridization we washed samples 2 × 20 minutes at 37°C using buffer containing 20% formamide and 2X SSC then rinsed the sample with 2X SSC. We then performed the copper(I)-catalyzed azide-alkyne cycloaddition (*“*click” reaction) by adding a solution containing 150 μM BTTAA, 75 μM copper sulfate, 2.5 mM L-ascorbic acid and 0.1% Triton-X 100 in 2X SSC to each sample and incubating at 37°C for 15-20 minutes. To prepare this solution, we first combined the BTTAA and copper sulfate, add the 2X SSC containing 0.1% Triton-X, and lastly add freshly dissolved L-ascorbic acid (19-20 mg of L-ascorbic acid sodium salt dissolved in 1 mL nuclease-free water). Once the L-ascorbic acid is added, we immediately added the solution to our samples. Following the click reaction, we rinsed samples once with 2X SSC then washed 1 × 20 minutes at 37°C with buffer containing 40% formamide in 2X SSC. After this wash, we performed the third round of hybridization with P9B and P9C amplifier probes in the amplifier hybridization buffer, followed by washes, click and post-click wash as described above. We continued with additional amplifier hybridizations (iterating between using MM2B+MM2C amplifier probes on even rounds and P9B+P9C amplifier probes on odd rounds) and washes, performing the click reaction during every odd round (3, 5, 7…).

After the post-click wash for round 7 or round 9, we added RNA FISH hybridization buffer (10% formamide and 10% dextran sulfate in 2X SSC) containing probes targeting P9B and P9C and coupled to Alexa Fluor 594 and Atto647n, respectively (see Supplementary Table 4 for sequences). We hybridized these probes overnight in humidified containers at 37°C then washed samples 2 × 30 minutes with washing buffer (10% formamide, 2X SSC) at 37°C, adding DAPI to the second wash to stain the nuclei. After these washes, we rinsed samples once with 2X SSC then replaced the 2X SSC and proceeded with imaging. To remove ClampFISH signal, we stripped dye-coupled probes as described above for RNA FISH.

### Immunofluorescence

We performed immunofluorescence using primary antibodies targeting total ERK (L34F12 Cell Signaling #4696) and phosphorylated ERK (p44/p42 ERK D12.14.4E Cell Signaling #4370). First, we rinsed cells 3 times with 5% BSA in PBS (5% BSA-PBS) then incubated at room temperature for 2 hours in 5% BSA-PBS containing 1:100 total ERK and 1:200 pERK antibodies. Next, we washed the cells 5 × 5 minutes with 5% BSA-PBS then incubated the cells at room temperature for 1 hour in 5% BSA-PBS containing 1:500 donkey anti-mouse secondary antibody conjugated to Cy3 (Jackson 715-165-150) and 1:500 goat anti-rabbit secondary antibody conjugated to Alexa Fluor 594 (Cell Signaling #8889). After the secondary incubation, we washed the cells 5 × 5 minutes with 5% BSA-PBS containing 50 ng/mL DAPI, then replaced the wash buffer with 2X SSC and proceeded with imaging as described below.

### RNA FISH and Immunofluorescence Imaging

We imaged RNA FISH samples on an inverted Nikon TI-E microscope equipped with a SOLA SE U-nIR light engine (Lumencor), an ORCA-Flash 4.0 V3 sCMOS camera (Hamamatsu), 20X Plan-Apo λ (Nikon MRD00205), 40X Plan-Fluor (MRH00401) and 60X Plan-Apo λ (MRD01605) objectives, and filter sets for DAPI, Cy3, Alexa Fluor 594, and Atto647N. For barcode ClampFISH and barcode HCR, we first acquired tiled images in a single Z-plane (scan) at 20X or 40X magnification, then, after identifying positions containing cells positive for resistant barcodes, we returned to those positions to acquire a Z-stack at 60X magnification. For subsequent rounds of single-molecule RNA FISH and ERK immunofluorescence, we acquired Z-stacks at 60X magnification. For scans, we used a Nikon Perfect Focus system to maintain focus across the imaging area.

### Image analysis

To identify Barcode RNA FISH-positive and GFP-positive cells in Supplementary Fig. 1f-h, we used custom MATLAB scripts to first stitch together scanned images, then identify individual cells using the DAPI nuclear signal. Next, we used a custom graphical user interface (GUI) to zoom in on the stitched image, view the barcode RNA FISH (Alexa647) signal, and interactively select barcode RNA FISH positive cells. After selecting all barcode RNA FISH-positive cells, we repeated the same process with GFP signal to select all GFP-positive cells without knowledge of the cells’ barcode RNA FISH status. We then extracted the spatial coordinates, barcode RNA FISH status, and GFP status for all cells, and plotted the results using custom R scripts available on Dropbox at https://www.dropbox.com/sh/u4sibi0fgorzk0p/AACmLLvqf0iY9GlZBzzuVbtTa?dl=0. MATLAB scripts for stitching scans and the custom GUI are available at https://github.com/arjunrajlaboratory/timemachineimageanalysis.

To identify Barcode RNA FISH positive cells for Rewind, we used custom MATLAB scripts to stitch, contrast and compress scan images (scripts available at https://github.com/arjunrajlaboratory/timemachineimageanalysis) then manually reviewed these stitched images. This review yielded positions containing candidate Barcode RNA FISH positive cells which we then re-imaged for verification at 60X magnification in multiple Z-planes. If we were uncertain about the fluorescence signal in a candidate cell (e.g. abnormal localization pattern, non-specific signal in multiple channels), we excluded the cell from imaging during subsequent rounds of RNA FISH or immunofluorescence.

For quantification of RNA FISH images we used custom MATLAB software available at: https://github.com/arjunrajlaboratory/rajlabimagetools. Briefly, the image analysis pipeline includes manual segmentation of cell boundaries, thresholding of each fluorescence channel in each cell to identify individual RNA FISH spots, and then extraction of spot counts for all channels and cells. After extracting spot counts, we analyzed RNA levels across single cells using custom R scripts available at https://www.dropbox.com/sh/u4sibi0fgorzk0p/AACmLLvqf0iY9GlZBzzuVbtTa?dl=0. In all figures, boxplots indicate the 25th, 50th and 75th percentiles with whiskers extending to 1.5 times the interquartile range. Notably, for some markers, we were not able to quantify expression in a few cells because of grossly abnormal or non-specific fluorescence signal (i.e. schmutz) or because we lost a cell during sequential hybridizations. We excluded data from these cells from analyses and as a result, some plots may contain slightly different numbers of points for different markers. For analyses involving dimensionality reduction (UMAP) or clustering, we only included cells with data for all assayed markers.

For the UMAP visualizations we used the Seurat v3.2.0 package (the versions of all dependent packages are documented in the plotting scripts on Dropbox and at https://www.dropbox.com/s/v66v41zryogmd78/RsessionInfo.txt?dl=0) ^40,41^. For the analysis shown in Fig. 4, we ran the UMAP algorithm on scaled RNA FISH data using the first 5 principal components and setting n_neighbors = 30 and min_dist = 0.3 (default settings). For the analyses shown in Fig. 5 and Fig. 6, we used the first 6 principal components and set min_dist = 0.6 to better visualize the number of cells expressing high levels of *DEPTOR*.

We adapted the RajLabImagetools pipeline for quantifying immunofluorescence images. After manually segmenting cells, we used custom MATLAB scripts to average fluorescence intensity within cell boundaries for each channel then took the maximum average fluorescence intensity across Z-planes. We additionally used DAPI signal to automate nuclei segmentation and separately quantified cytoplasmic and nuclear immunofluorescence intensity. We found qualitatively similar results for both cytoplasmic and nuclear ERK immunofluorescence quantification (Supplementary Fig. 8).

For quantification of cell and colony numbers following vemurafenib treatment, we used custom MATLAB software available at https://github.com/arjunrajlaboratory/colonycounting_v2. The analysis pipeline involves stitching the tiled DAPI images, manually segmenting individual wells and colonies, identifying individual cells based on DAPI signal, and then extraction of cell counts from the entire well and each colony. We analyzed the extracted cell counts using custom R scripts available at: https://www.dropbox.com/sh/u4sibi0fgorzk0p/AACmLLvqf0iY9GlZBzzuVbtTa?dl=0. We used a separate MATLAB script (https://www.dropbox.com/s/xnwtmw8rh8ec3ij/countCellsTimeMachineScans.m?dl=0) to quantify the number of cells imaged in our Carbon Copies.

To assign individual primed cells (marked by barcode RNA FISH signal) to subclones (Supplementary Fig. 10), we first extracted the spatial position of each image in the whole-well scans containing at least 1 primed cell. We then calculated the Euclidean distance between these images and used these distances to perform hierarchical clustering. Visual inspection of the clustering revealed a clear distance threshold of < 2mm for grouping subclones of closely related (and therefore neighboring) primed cells; thus, all primed cells within these groups were assigned to belong to the same subclone. To further check our subclone assignments, we manually inspected all barcode RNA FISH images and found that primed cells assigned to the same subclone had similar barcode RNA FISH signal intensity and intracellular patterns, while this signal similarity was not observed for primed cells assigned to different subclones. Most primed cells from different subclones were at least 7 mm apart, and for the few cases of primed cells located between 2 mm − 7 mm apart, we observed that these cells had distinct barcode RNA FISH signal patterns consistent with them belonging to separate subclones. This clear spatial separation gave us confidence in our ability to accurately assign individual cells to particular subclones.

### RNA Sequencing and Analyses

We extracted RNA from fixed cells after barcode RNA FISH and sorting using the NucleoSpin total RNA FFPE XS kit (Takara). We performed cell lysis and reverse cross-linking at 50°C for 90 minutes and otherwise followed the manufacturer’s protocol. After RNA extraction, we prepared sequencing libraries using the NEBNext single-cell/low-input RNA sequencing library preparation kit for Illumina (NEB) then performed paired-end sequencing of these libraries (38 cycles read 1 + 37 cycles read 2) on a NextSeq 500 (Illumina). After sequencing, we aligned reads to the human genome (assembly 19; hg19) using STAR^42^ v2.5.2a and counted uniquely mapped reads with HTSeq^43^ v0.6.1.

We performed differential expression analysis in R v3.6.3 using DESeq2^44^ v1.22.2 and with data from at least 2 biological replicates for each sample and condition. Biological replicates were sorted on separate days using distinct barcode RNA FISH probe sets. We considered a gene to be differentially expressed if the comparison between 2 conditions yielded a log_2_ fold change of ≥1 or ≤ −1 and adjusted p-value of ≤0.1. For determining candidate markers for primed cells requiring DOT1L inhibition (Fig. 5) we compared primed and non-primed subpopulations sorted from both DOT1L inhibitor and vehicle control Carbon Copies and modelled the biological replicate and DOT1L inhibitor treatment as covariates in the design formula for DESeq2. We chose to include data from both DMSO- and DOT1L-inhibitor-treated Carbon Copies (2 replicates each) in the analysis and model DOT1L inhibitor treatment as a covariate due to the modest effects of DOT1L inhibitor treatment alone on gene expression (Fig. 6e,f, Supplementary Fig. 14d,e, and Supplementary Fig. 16c,d) and our particular interest in identifying gene expression markers that distinguish various subpopulations of primed cells. We performed hierarchical clustering and principal component analysis on log_2_ transformed TPM values using R v3.6.3.

We tested for enrichment of differentially expressed genes among gene ontologies and pathways (KEGG, REACTOME, WikiPathway) using WebGestaltR. If a differentially expressed gene was included in one or more enriched GO term or pathway, we chose a consensus annotation (e.g. ECM organization and cell migration) for that gene. Otherwise, we attempted to assign a gene annotation by manual review. Our resulting gene annotation can be found in Supplementary Table 8.

### Reporting Summary

Further information on research design is available in the Nature Research Reporting Summary linked to this article.

## Data Availability

All RNA sequencing data generated for this study are available on GEO (accession #GSE161300). Additional sequencing and imaging data are available on Dropbox at https://www.dropbox.com/sh/mmeg3mckrpridu3/AAALBaMLoJsJiQC2-lrVY0Cva?dl=0 and upon request of the corresponding author.

## Code Availability

Software used to segment cells and quantify RNA spots is available at https://github.com/arjunrajlaboratory/rajlabimagetools. Software used to stitch, segment and quantify scan images of resistant colonies is available at https://github.com/arjunrajlaboratory/colonycounting_v2. Additional custom image analysis scripts are available at https://github.com/arjunrajlaboratory/timemachineimageanalysis. The pipeline used for barcode sequencing analysis is available at https://github.com/arjunrajlaboratory/timemachine.

