## Supplemental Figures 1-16 for "Variability within rare cell states enables multiple paths towards drug resistance"

### Supplementary Figure 1

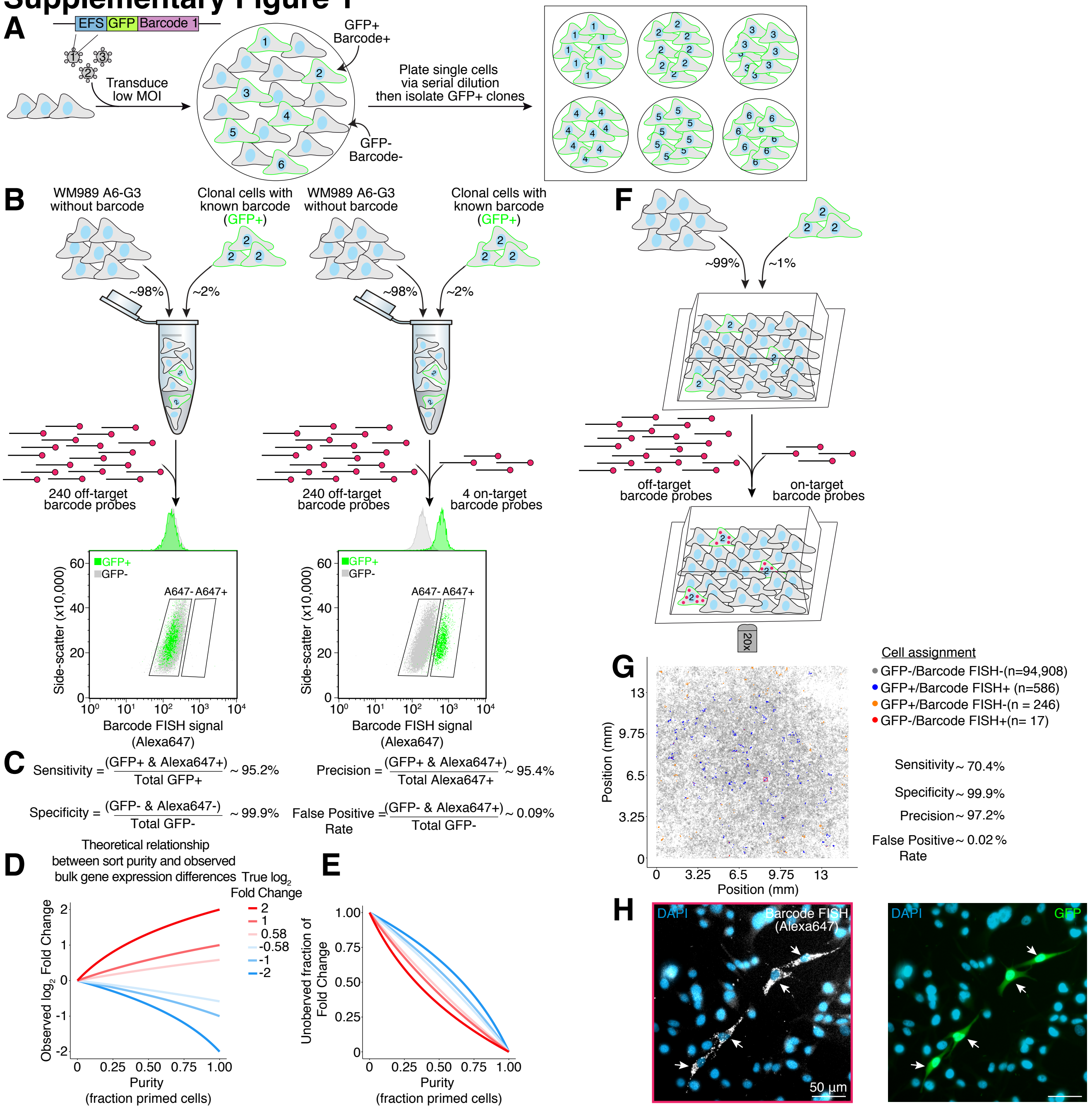


**Supplementary Figure 1. Detection and isolation of cells expressing unique Rewind barcodes using RNA FISH. A.** The Rewind construct encodes a 100 nucleotide barcode sequence ("WSN" repeated) in the 3' UTR of *GFP* downstream of a truncated *EEF1A1* promoter (EFS). For use in optimizing barcode RNA FISH, we derived clonal cell lines each expressing a single barcode and identified the barcode sequence in each cell line via Sanger sequencing. **B-C.** We combined GFP- WM989 A6-G3 cells (~98% of cells) with a GFP+ WM989 A6-G3 subclone carrying a single, known Rewind barcode (~2% of cells), then performed barcode RNA FISH (HCR protocol) using 4 probes targeting the known barcode along with 240 "off-target" probes (designed to target barcodes sequenced in other experiments). In parallel, we hybridized a second mix of cells with only the off-target probes. After hybridization, we ran these samples on a FACS instrument and used GFP fluorescence as ground truth for estimating the sensitivity and specificity of the barcode RNA FISH signal (Alexa647). To isolate rarer subpopulations with Rewind (e.g. primed cells from a Carbon Copy), we used a more conservative Alexa647 gate in an attempt to further minimize false positives. **D.** Contamination of rare subpopulations isolated by FACS with off-target cells can dilute expression changes in bulk RNA-seq analyses. We computed the theoretical observed $\log_2$ fold change for primed cells vs. non-primed cells across a range of sample purity levels (x-axis) and actual $\log_2$ fold change values (line colors). **E.** We calculated the theoretical difference in observed vs. actual fold change (normalized to actual fold change; y-axis) across a range of sample purity levels (x-axis). Of note, the larger differences for negative fold change values (blue lines) suggest that bulk analyses of enriched primed cells may be less able to detect decreased levels of gene expression depending on the degree of enrichment. **F.** To assess the accuracy of Rewind for imaging-based applications, we mixed together and plated GFP- WM989 A6-G3 cells (~99% of cells) with a GFP+ WM989 A6-G3 subclone carrying a single, known Rewind barcode (~1% of cells). We cultured the cells for 4 days before fixing the cells and performing barcode RNA FISH (HCR protocol) using 4 probes targeting the known barcode and 120 probes targeting random off-target barcodes. We then imaged the entire well with a tiled scan at 20X magnification and identified all barcode FISH+ cells (Alexa647+) followed by all GFP+ cells (true positives) using semi-automated software (see Methods). **G.** We plotted the position of each cell analyzed in F and color-coded each point by its assigned GFP and barcode RNA FISH status. The red square corresponds to the position of the micrographs shown in G. The number of cells with each label are indicated in parentheses in the legend. **H.** Example 20X micrographs showing barcode RNA FISH signal (Alexa647; left) and GFP signal (right) overlaid with DAPI signal. White arrows point to 4 GFP+/Alexa647+ cells out of 586 similar cells identified in the well (n = 1 biological replicate).

### Supplementary Figure 2

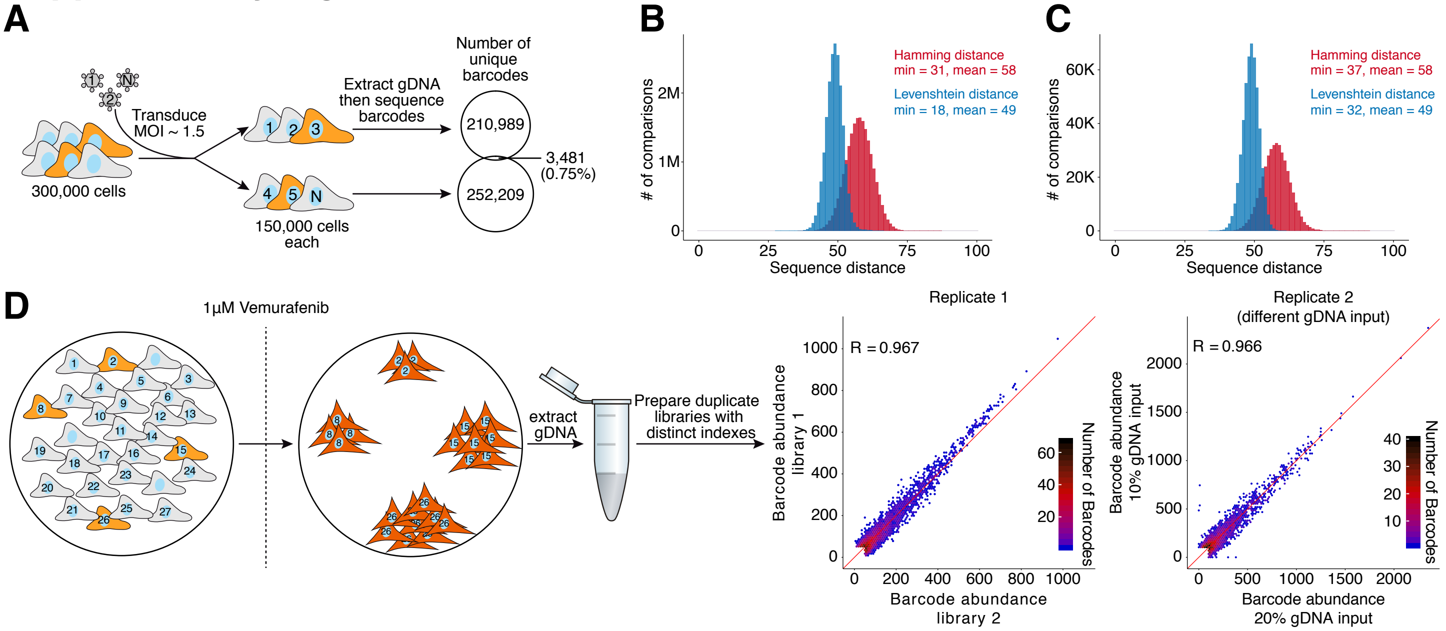


**Supplementary Figure 2. The Rewind barcode library can uniquely label 100,000s of cells with transcribed barcodes that can be identified via sequencing. A.** Critical for Rewind is the ability to uniquely label enough cells with the transcribed barcodes to observe rare phenomena such as drug resistance (frequency < 1:1000). To test this empirically, we separately transduced 2 groups of 150,000 cells at an MOI of ~1.5 , cultured the cells for ~1 day then extracted genomic DNA (gDNA) and sequenced their barcodes. Consistent with the starting cell number and MOI, we observed between 210,000 and 253,000 barcodes in the two samples ($\geq 2$ unique reads) with fewer than 3,500 (~0.75%) shared between the two groups. **B.** To estimate the sequence complexity of Rewind barcodes, we calculated pairwise sequence distances for 20 million randomly sampled pairs of barcodes identified in A. Plotted is the distribution of pairwise Hamming (red) and Levenshtein (blue) distances. Of note, to reduce artifacts from library preparation and sequencing, we collapsed extremely similar barcodes (Levenshtein distance $\leq 8$ for this plot; see Methods for details). We observed similar pairwise distances using different thresholds for collapsing similar barcodes (for example, panel C uses a Levenshtein distance threshold of $\leq 4$). **C.** To estimate barcode complexity in cells following selection with vemurafenib, we analyzed barcode sequencing data from barcoded WM989 A6-G3 (~200,000 cells transduced at MOI ~ 1.5) after 3 weeks of treatment with 1 μM vemurafenib. Plotted is distribution of observed Hamming (red) and Levenshtein (blue) distances for all pairwise comparisons (~500,000) of the 1,000 barcodes with the highest read counts (corresponding to the most resistant clones). **D.** We tested the reproducibility of our barcode sequencing protocol by preparing separate libraries with unique indexes using the same starting gDNA. As shown in the scatter plots we see a high correspondence in barcode abundances (UMIs per million; see Methods for description of barcode count normalization) between these replicate libraries, even when using different amounts of gDNA (2 independent experiments shown). Plotted are all barcode sequences with at least 50 reads per million in at least one of the two samples. We believe these data also suggest that our barcode sequencing protocol is quantitative, however we acknowledge the possibility that differences between barcode sequences could systematically bias library preparation. Our validation of barcode RNA FISH probes designed to target more abundant versus less abundant resistant cells (Figure 4) further suggests that our sequencing data provides a quantitative estimate of clone abundance.

### Supplementary Figure 3

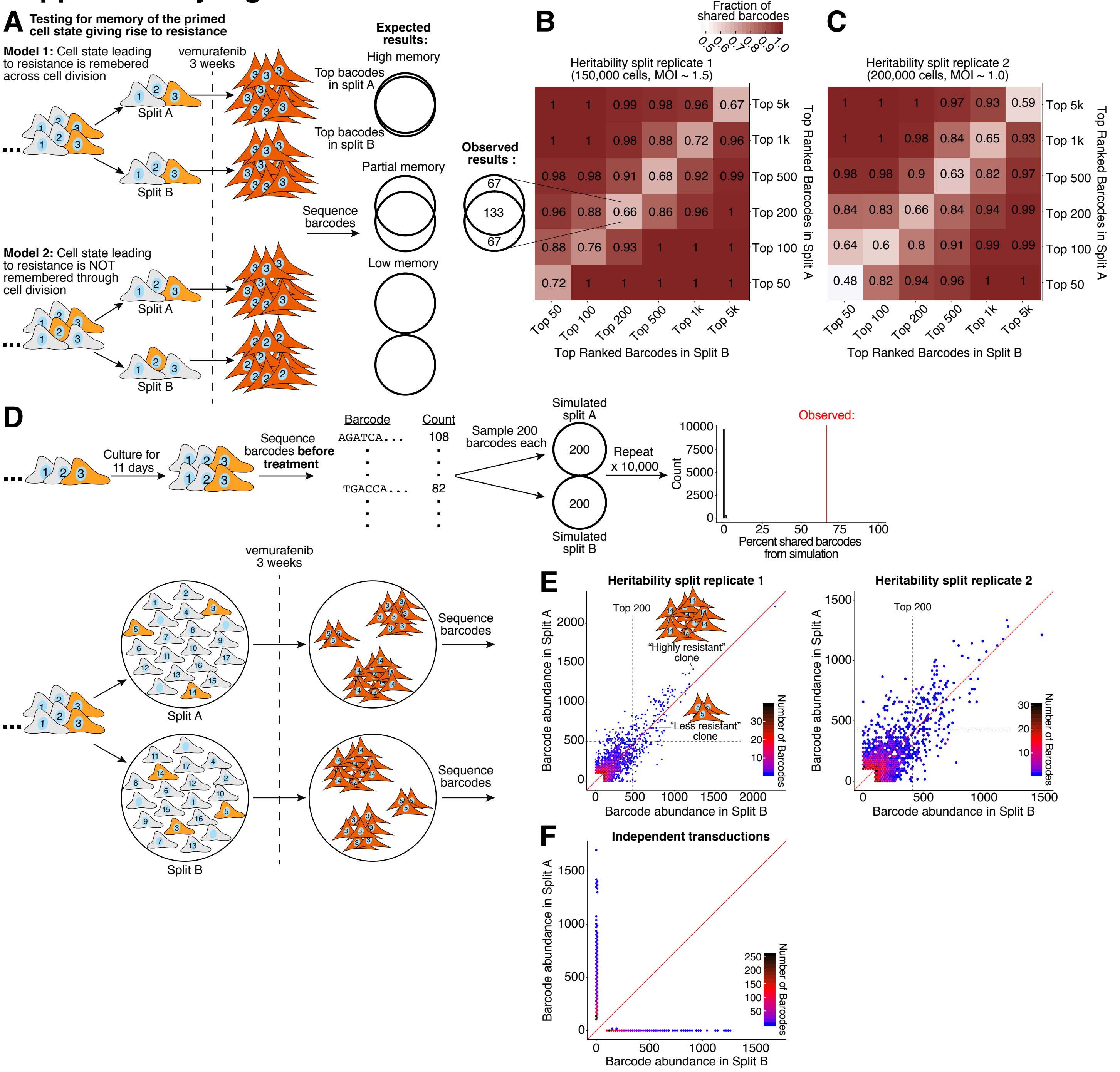


**Supplementary Figure 3. Barcode sequencing of "twin" cultures treated with vemurafenib suggests that the primed cell state is maintained through several cell divisions. A.** Schematic of the cellular barcoding approach used to test whether the primed cell state is "remembered" through cell division. We transduced ~150,000-200,000 WM989 A6-G3 cells with our Rewind barcode library and allowed the cells to divide for 11-12 days (~4-5 cell divisions). We then split the culture in two and treated both halves with 1 µM vemurafenib for 3 weeks. Finally we sequenced the barcodes in genomic DNA extracted from each culture then ranked the barcodes by abundance to identify those likely derived from resistant colonies (expected 100-400 unique barcodes from resistant colonies). In the absence of memory of the primed cell state, we expected to find unique sets of barcodes emerging in the two parallel cultures. In the presence of partial or complete memory, we expected to find some overlap in the barcodes identified in each culture. **B-C.** Heatmap shows the proportion of barcodes shared between the parallel cultures at different rank thresholds. For our Rewind experiments, we selected the top 100-200 barcodes for RNA FISH probe design. **D.** We wanted to rule out the possibility that differences in division rate between cells before adding vemurafenib could skew the distribution of barcodes enough to generate the observed barcode overlap by chance alone. We therefore sequenced barcoded cells after 11 days of growth (before vemurafenib treatment) to estimate the change in the barcode distribution due to differences in cell growth. We then simulated the split and vemurafenib treatment in A by randomly sampling 2 groups of 200 cells each from the observed barcode distribution and calculating the proportion of shared barcodes. The histogram shows the results of repeating this simulation 10,000 times (gray bars) with the red line indicating the experimentally observed proportion of shared barcodes (from B). **E.** We compared the abundance of barcodes from parallel cultures in A-C by plotting all barcodes with at least 100 UMIs per million in at least one sample (see Methods for description of normalization).To better visualize lower abundance barcodes, we binned the barcodes by count and colored each bin by its number of unique barcodes. Based on the observed correlation in barcode abundance between parallel cultures, we reasoned that differences in the number of cells that make up a vemurafenib resistant clone (which varies by more than an order of magnitude) are at least partially pre-determined in the initial primed population 3 weeks earlier. **F.** Reassuringly, we do not observe a correlation in barcode counts between vemurafenib resistant cells from independent transductions

### Supplementary Figure 4

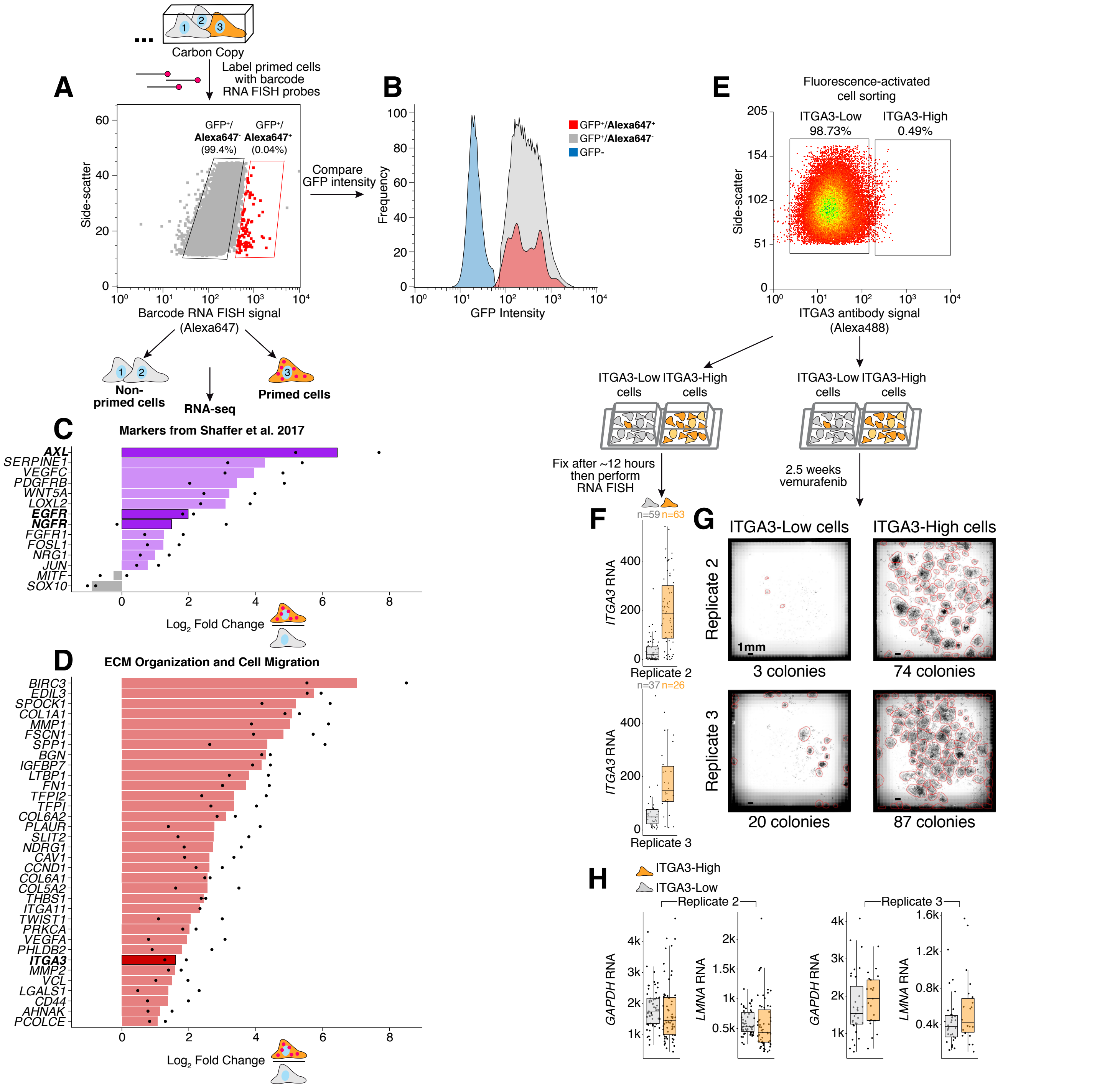

**Supplementary Figure 4. RNA sequencing of primed WM989 A6-G3 isolated using Rewind identifies *ITGA3* as a prospective marker of vemurafenib resistance. A.** FACS data for the Rewind experiment presented in Figure 1. **B.** We observed similar levels of GFP fluorescence in sorted primed (GFP+/Alexa647+) and non-primed (GFP+/Alexa647-) cells in A. **C.** We compared the expression of genes previously implicated in vemurafenib resistance (Shaffer et al. 2017) in primed and non-primed cells isolated using Rewind. Bargraph indicates the average $log_2$ fold change in expression in primed versus non-primed cells with individual replicates indicated as points. We previously demonstrated that cells expressing the genes in bold (*AXL*, *EGFR*, and *NGFR*) are more likely to form vemurafenib resistant colonies and these same cells express lower levels of *SOX10* and *MITF* (gray bars) (Shaffer et al. 2017). **D.** We found an enrichment for genes associated with ECM organization and cell migration among differentially expressed genes comparing primed cells to non-primed cells (see Methods for pathway analyses and Supplementary Table 6 for FDR values). Bargraph shows the average $log_2$ fold change in expression in primed versus non-primed cells with individual replicates indicated as points. We validated *ITGA3* (bolded) as a predictive marker of vemurafenib resistance (Figure 1 and panels E-G). We did not detect expression of *ITGA11* in non-primed cells in 1 of 2 replicates (it was detected in primed cells) and the presented data corresponds to the $log_2$ fold change for 1 replicate. **E.** We stained cells with an anti-ITGA3 antibody, then sorted equal numbers of the brightest ~0.5% (ITGA3-High) and remaining ~99% (ITGA3-Low) cells. We plated ~1/3 of these cells onto 1 plate for measuring *ITGA3* expression by RNA FISH and plated the rest onto a separate plate for treatment with 1 µM vemurafenib. After ~18 days of treatment, we fixed the cells, stained nuclei with DAPI then imaged the wells to quantify the number of resistant colonies and cells. **F.** Quantification of *ITGA3* RNA by RNA FISH in sorted cells from E. Each point corresponds to an individual cell with the total number of cells indicated above each boxplot. **G.** Whole-well scans of sorted cells from E after vemurafenib treatment. Shown are stitched micrographs from 2 of 3 biological replicates (3rd replicate shown in Figure 1). **H.** By RNA FISH, we observed similar levels of *GAPDH* and *LMNA* RNA in ITGA3-High and ITGA3-Low cells from E. For all boxplots, center line indicates median, box limits indicate 25th and 75th percentiles, and whiskers extend to 1.5 times the interquartile range.

### Supplementary Figure 5

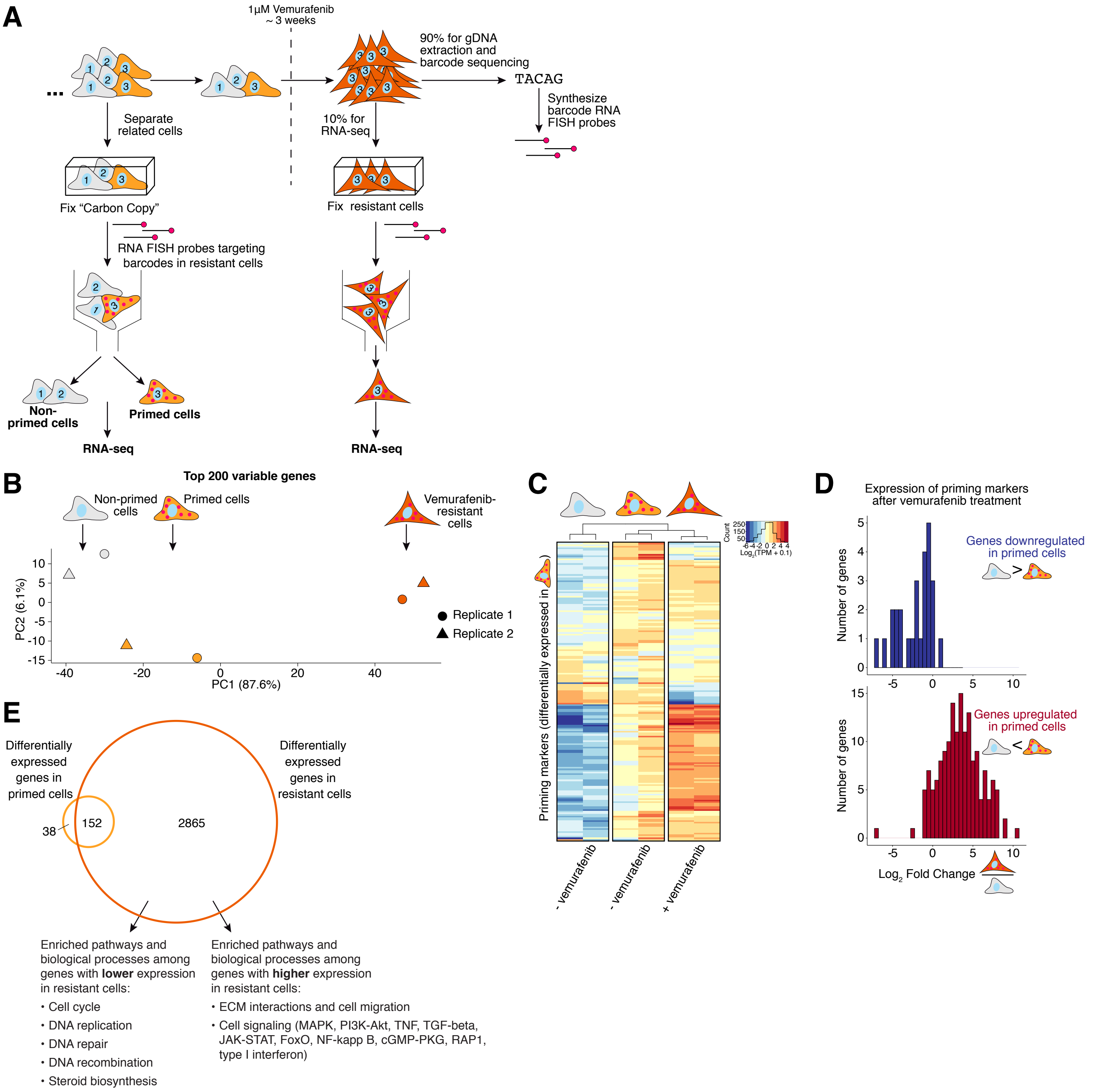


**Supplementary Figure 5. Most priming markers remain transcriptionally altered after 3 weeks of vemurafenib treatment, and an additional ~3000 genes become differentially expressed. A.** When performing Rewind in WM989 A6-G3 cells, we collected ~10% of resistant cells for RNA sequencing alongside cells sorted from our Carbon Copy (see Methods for details). Profiling these cells enabled us to ask how gene expression changed in primed cells after vemurafenib treatment during their transition into resistant cells. **B.** We performed principal component analysis using $\log_2$(TPM+0.1) values for the 200 protein-coding genes with the largest variance across samples. Shown are the position of each sample along the first 2 principal components. Notably, primed cells are positioned much closer to non-primed cells than to resistant cells along the first principal component (which explains ~88% of the variance). **C.** Heatmap shows the expression ($\log_2$(TPM+0.1)) of priming markers in non-primed cells (left), primed cells (middle) and vemurafenib resistant cells (right). We defined priming markers as genes differentially expressed (p-adjusted ≤ 0.1 and abs($\log_2$ fold change) ≥1) in primed versus non-primed cells. **D.** Histogram shows the $\log_2$ fold change in expression in resistant cells for priming markers downregulated in primed cells (top, blue) or upregulated in primed cells (bottom, red). Of note, the majority of gene expression changes observed in primed cells persist during their transition into resistant cells following vemurafenib treatment. **E.** We found more than 3,000 genes differentially expressed comparing resistant cells to drug-naive non-primed cells of which the ~200 priming markers represented a small subset. We performed pathway and gene-ontology analyses on the genes differentially expressed only in resistant cells and highlight several recurring annotations (see Supplementary Table 7 for a complete list and FDR values).

### Supplementary Figure 6

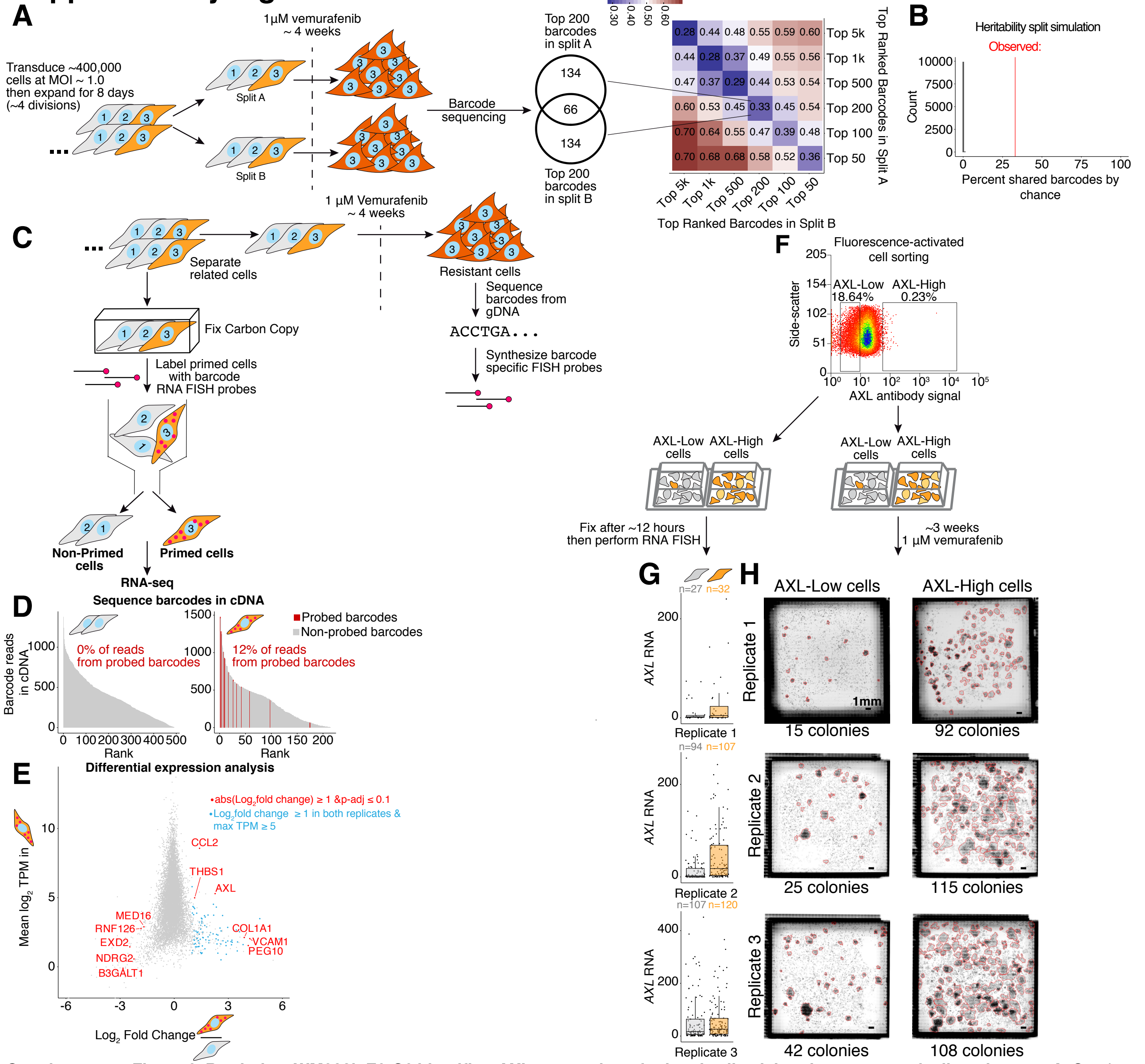


**Supplementary Figure 6. Rewind on WM983b E9-C6 identifies *AXL* as a marker of primed cells giving rise to vemurafenib resistance. A.** Starting with ~400,000 cells transduced at an MOI ~ 1.0, we repeated the "heritability-split" experiment in WM983b E9-C6 cells to determine if the initial primed state giving rise to vemurafenib resistance was maintained through several cell divisions. The Venn diagram and heatmap show the proportion of barcodes shared between the parallel cultures at different rank thresholds. **B.** To estimate the probability of seeing the observed fraction of shared barcodes by chance, we simulated the experiment using data from WM93b E9-C6 cells transduced as in A then cultured for 7 days before sequencing. We simulated the split and vemurafenib treatment by randomly sampling 2 groups of 200 cells each from the observed barcode distribution then calculated the proportion of shared barcodes. The histogram shows the results of repeating this simulation 10,000 times (gray bars) with the red line indicating the observed proportion of shared barcodes from A. **C.** Using Rewind we isolated primed WM983b E9-C6 cells from Carbon Copies fixed prior to vemurafenib treatment (n = 2 biological replicates). We then performed RNA sequencing and barcode sequencing on cDNA prepared from sorted cells. We expanded barcoded cells for ~4 population doublings before dividing the cells into the Carbon Copy or the drug-treated half. **D.** Bargraphs show the abundance (y-axis) and rank (x-axis) of barcodes sequenced from cDNA (≥ 5 normalized reads). Red bars correspond to barcodes targeted by our probe set and gray bars correspond to non-targeted barcodes. Inset shows the percent of barcode sequencing reads that match a probe-targeted barcode. Of note, the baseline frequency of on-target barcodes was ~0.015% (60/400,000). **E.** We used DESeq2 to identify differentially expressed genes (p-adjusted ≤ 0.1 and abs($log_2$ fold change) ≥1) in primed cells versus non-primed cells (red points). Compared to similar experiments in WM989 A6-G3, fewer genes passed our significance cutoff which may reflect the shorter memory of the primed cells state (see A) and the lower purity of our Rewind sort (D). We highlighted in blue genes that did not pass our significance cutoff but showed ≥ 2-fold higher expression in primed cells in 2 out of 2 replicates. **F.** We sorted drug-naive cells expressing high levels of *AXL* and low levels of *AXL* then compared their response to vemurafenib treatment. **G.** With ~ 1/3 of the cells, we performed RNA FISH to measure *AXL* expression in the 2 sorted populations. Points indicate the levels of *AXL* RNA in individual cells with the total number of cells indicated above each plot. For all boxplots, center line indicates median, box limits indicate 25h and 75th percentiles, and whiskers extend to 1.5 times the interquartile range. **H.** We treated the remaining sorted cells with vemurafenib for 3 weeks then imaged the wells to quantify the number of resistant colonies. Shown are stitched, whole-well scans of sorted cells from F after vemurafenib treatment (n = 3 biological replicates).

### Supplementary Figure 7

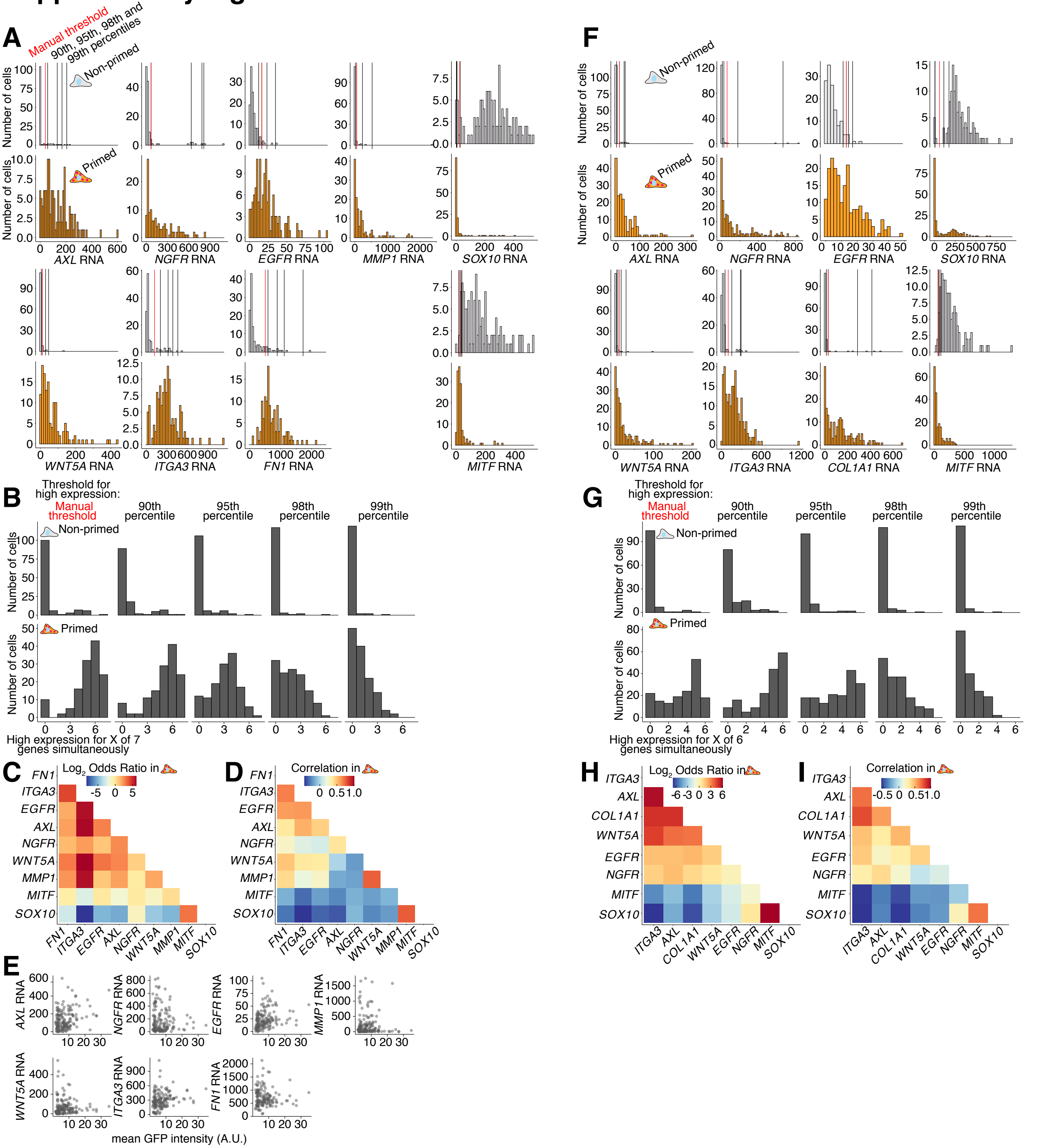

**Supplementary Figure 7. Primed WM989 A6-G3 cells express high levels of multiple markers simultaneously. A.** As in Figure 2, we identified primed cells that give rise to vemurafenib resistance using Rewind then measured single-cell expression of 7 priming markers, *SOX10* and *MITF* using RNA FISH. Histograms show the expression distributions for these genes in primed cells (orange) and randomly selected non-primed cells (gray). We used these distributions to set thresholds for binning high expressing cells and, in turn, characterizing the co-expression of these markers in single cells. The black vertical lines correspond to the 90th, 95th, 98th and 99th percentiles of expression in non-primed cells and the red vertical line corresponds to the threshold used for Figure 2. **B.** For the indicated expression thresholds, histograms show the number of primed cells (bottom row) and non-primed cells (top row) that express high levels multiple markers simultaneously (number of markers indicated on X-axis). **C-D.** For all pairs of markers, we calculated odds ratios for co-expression (at levels greater than the manual thresholds from A) and Pearson correlation in gene expression in primed cells. **E.** Comparison of single-cell gene expression with GFP fluorescence intensity in primed cells suggests that expression of priming markers is not associated with higher overall expression capacity. **F-I.** We repeated the analyses in A-D on non-primed and primed cells from a separate experiment. These primed cells correspond to cells that do not require DOT1L inhibition to become vemurafenib resistant from a Carbon Copy treated with vehicle control (DMSO) for 5 days before fixation (data also used in Figure 5).

### Supplementary Figure 8

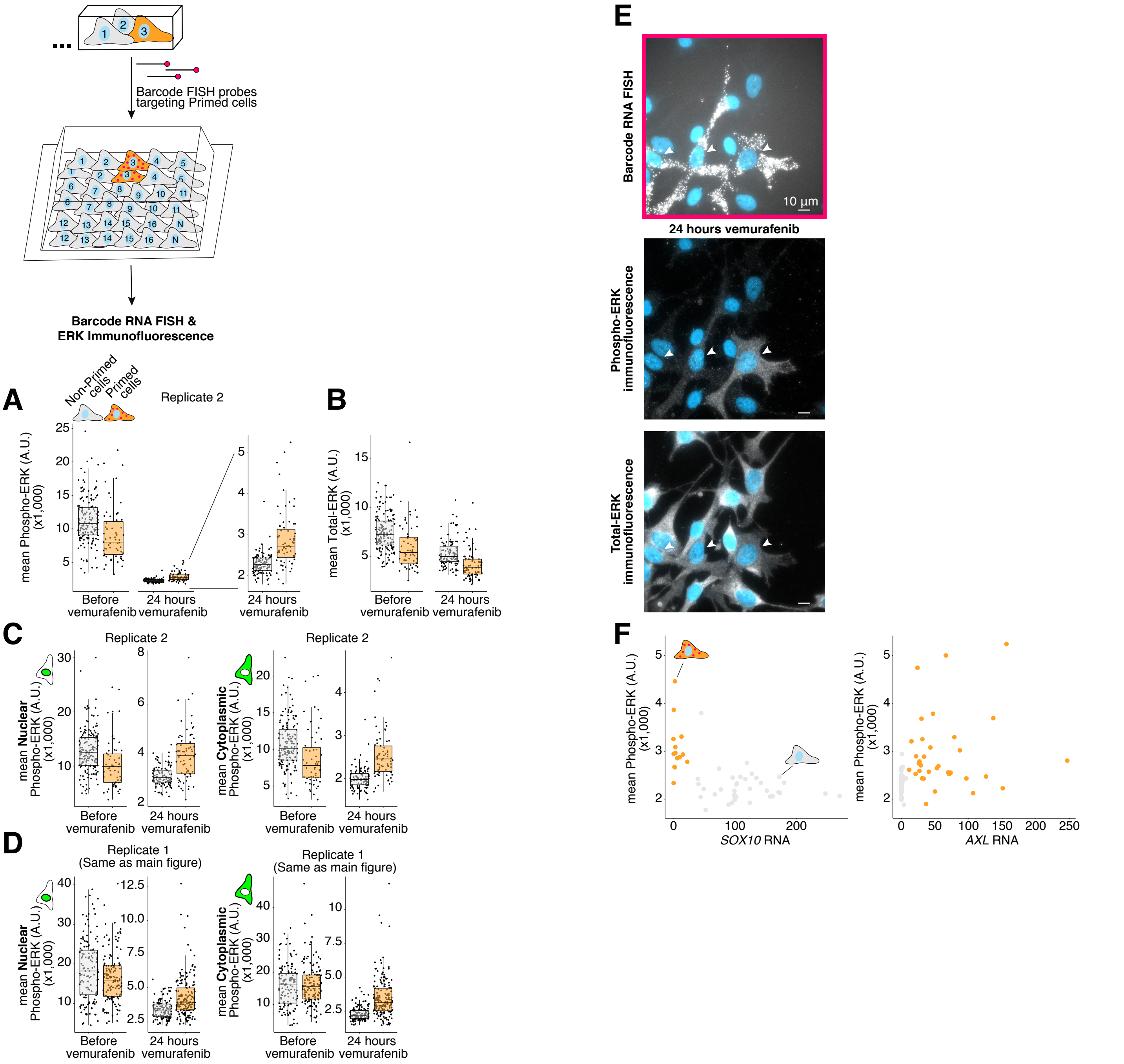


**Supplementary Figure 8. Primed cells giving rise to vemurafenib resistance have higher levels of phosphorylated ERK than non-primed cells 24 hours after vemurafenib treatment, but there remains cell to cell variability. A-B.** We used Rewind to quantify total ERK and dual-phospho ERK (p44/p42, pERK) levels in primed cells (orange) and non-primed cells (gray) before and 24 hours after vemurafenib treatment. Shown is a biological replicate of the experiment shown in Figure 3. For this second replicate, we amplified the barcode RNA FISH signal using ClampFISH instead of HCR (see Methods for details) and identified 130 primed cells (n = 59 before and n = 71 after vemurafenib treatment) and 271 non-primed cells (n = 162 before and n = 109 after vemurafenib treatment). Each point corresponds to an individual cell. **C-D.** In addition to average pERK across the entire cell, we calculated average pERK within just the cell nucleus (left) or just the cytoplasm (right). We found similar results using all of these metrics. For all boxplots (including panels A-B), center line indicates median, box limits indicate 25h and 75th percentiles, and whiskers extend to 1.5 times the interquartile range. **E.** While on average primed cells had higher levels of pERK, we observed cell-to-cell variability in several clusters of closely-related primed cells. Micrographs (60X) show one such example with arrowheads pointing towards primed cells (n = 2 biological replicates). We speculate that this variability may be a result of pulsatile MAPK signaling (which has been documented in other melanoma cell lines; Gerosa et al. 2019), and our snapshot measurement of ERK phosphorylation via immunofluorescence. **F.** After identifying primed cells in situ, we performed both single-molecule RNA FISH and immunofluorescence to measure gene expression and pERK levels in the same single cells. Shown is the relationship between pERK levels and *AXL* (left) or *SOX10* (right) expression in individual primed (orange points) and non-primed (gray points) cells. Within the primed cell population, we observe a fairly low correlation between pERK levels and expression of these markers, which we speculate may be a result of MAPK signalling fluctuating on a faster timescale than changes in gene expression.

### Supplementary Figure 9

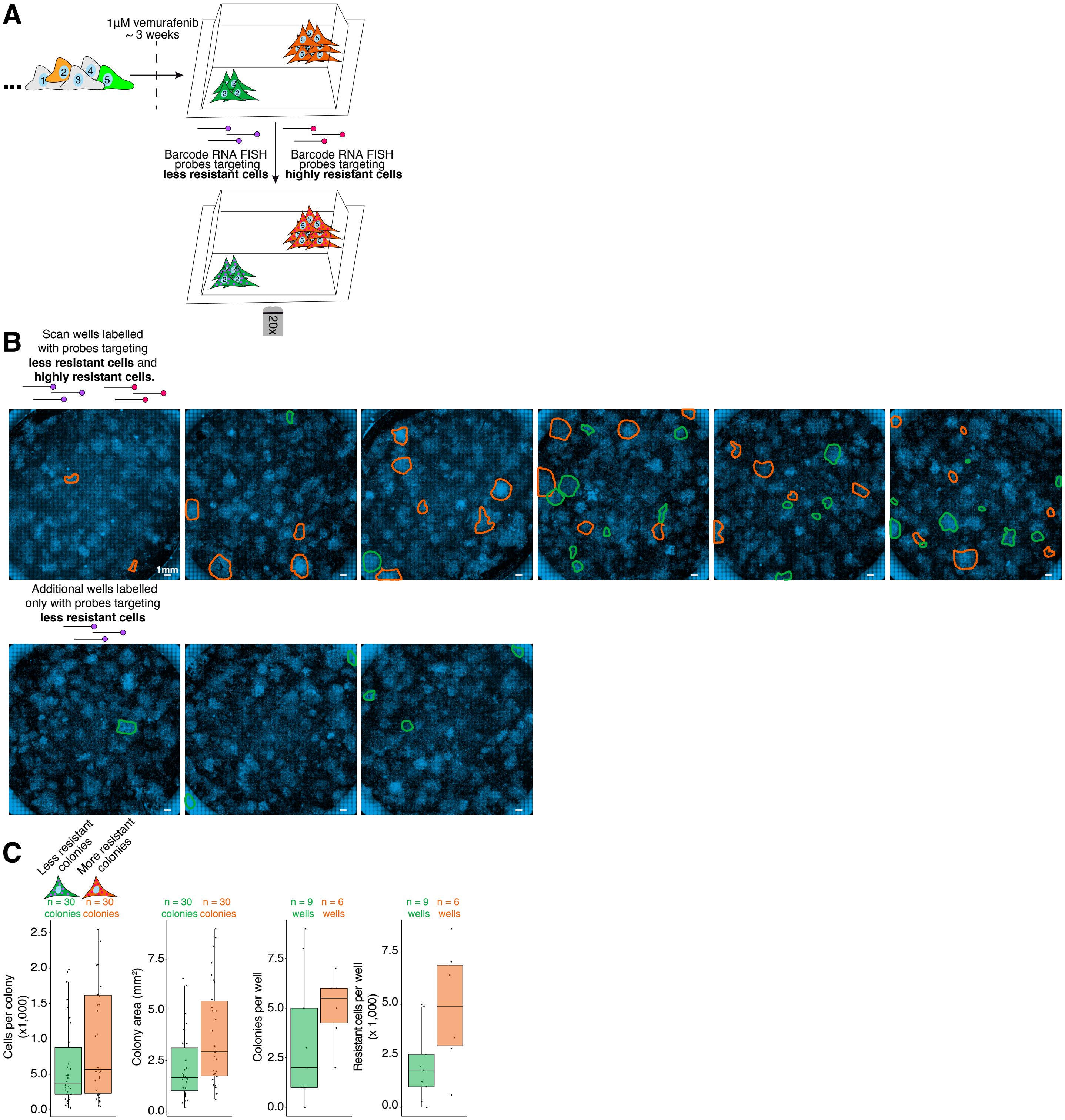


**Supplementary Figure 9. Barcode RNA FISH can distinguish highly resistant clones from less resistant clones. A.** As described in the Results for Figure 4, we sequenced barcodes from vemurafenib resistant cells then designed separate RNA FISH probe sets targeting 30 of the most abundant barcodes (ranked in the top ~ 50 most abundant) and 30 less abundant barcodes (ranked between ~50-100 by abundance). We reasoned that these two groups correspond to clones (cells sharing identical barcodes) with greater and fewer resistant cells (see Supplementary Figs. 2d and 3e for further reasoning). We refer to these two groups as “highly resistant” and “less resistant” as these groups roughly correspond to degrees of fitness in drug and our colleagues found the terms “more abundant” and “less abundant confusing. To empirically test that our probe sets distinguish different groups of resistant cells, we labeled resistant colonies derived from the same population of barcoded cells with our two probe sets, each coupled to a distinct fluorescent dye (Alexa Fluor 546 or Alexa Fluor 647). We then imaged the cells and quantified the number of colonies and resistant cells labeled with each probe set. **B.** Shown are stitched micrographs of scan images of barocde RNA FISH labelled resistant cells. We used custom software to annotate colonies labeled with our “highly resistant” (orange) or “less resistant” (green) probe sets (see Methods). To confirm that the observed differences in the number of cells labelled with each probe set was not due to differences in HCR hairpin and dye, we swapped the initiator sequence (and corresponding hairpin and dye) for the less resistant probe set, and labelled an additional 3 wells shown in the bottom row. (n = 1 biological replicate). **C.** Quantification of the number of colonies and number of cells labeled with each probe set. For all boxplots, center line indicates median, box limits indicate 25h and 75th percentiles and whiskers extend to 1.5 times the interquartile range.

#### Supplementary Figure 10

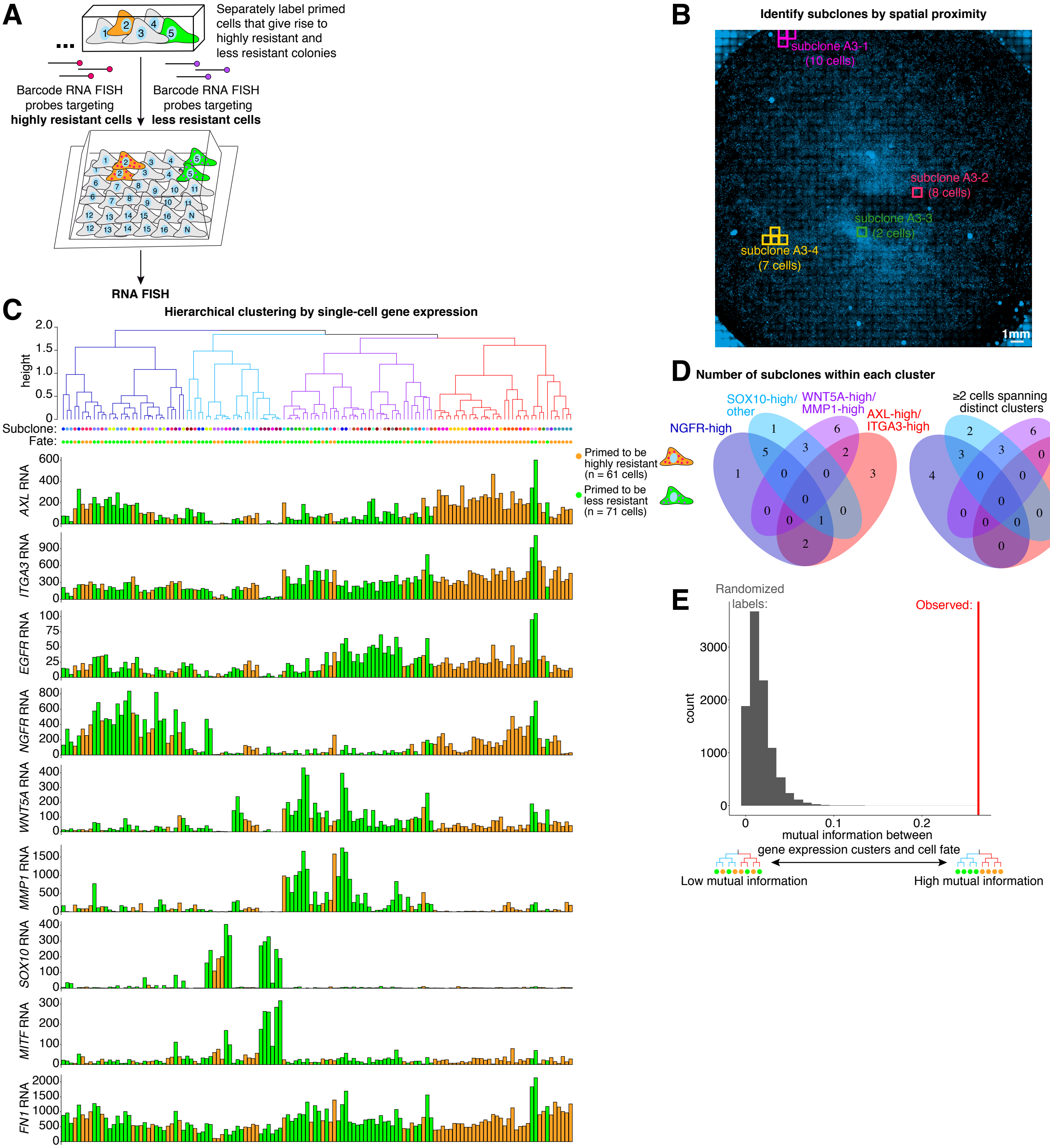


**Supplementary Figure 10. Rewind uncovers multiple, slowly-transitioning primed cell states. A.** We used Rewind to identify primed cells in situ, then measured expression of 9 genes in single cells using RNA FISH. As in Figure 4, we used separate barcode RNA FISH probe sets coupled to distinct fluorophores to distinguish primed cells giving rise highly-resistant or less-resistant colonies. **B.** Shown is a stitched micrograph of whole-well, 20X scan of a Carbon Copy with DAPI signal pseudocolored in blue. The colored squares outline image tiles containing at least 1 primed cell (barcode RNA FISH positive). To identify subclones of closely-related primed cells, we grouped primed cells located within a distance of 2 mm (see Methods for further details). The color of each square indicates a separate subclone with the number of primed cells within the subclone shown in parentheses. **C.** Dendrogram shows the results of hierarchical clustering of the single-cell gene expression data for primed cells. The branch colors indicate the 4 clusters of primed cells. Below the dendrogram, we used points to label the subclone (top row) and resistance fate (bottom row) of each primed cell (terminal branch). Below these points, we plotted the expression of each marker gene in each primed cell. The color of each bar indicates the resistance fate of the corresponding primed cell. **D.** Venn diagram indicates the number of subclones containing primed cells belonging to a single cluster from C (edges) or a mix of clusters (overlaps). For the Venn diagram on the far right, subclones were defined as mixed only if at least 2 primed cells were shared between two or more clusters. **E.** We calculated the mutual information between the gene-expression-cluster labels and resistance-fate labels for primed cells. We then permuted the cluster labels 10,000 times and recalculated the mutual information each time. Plotted is the distribution of mutual information values for the permuted data (gray bars) and the value for the observed data (red line).

### Supplementary Figure 11

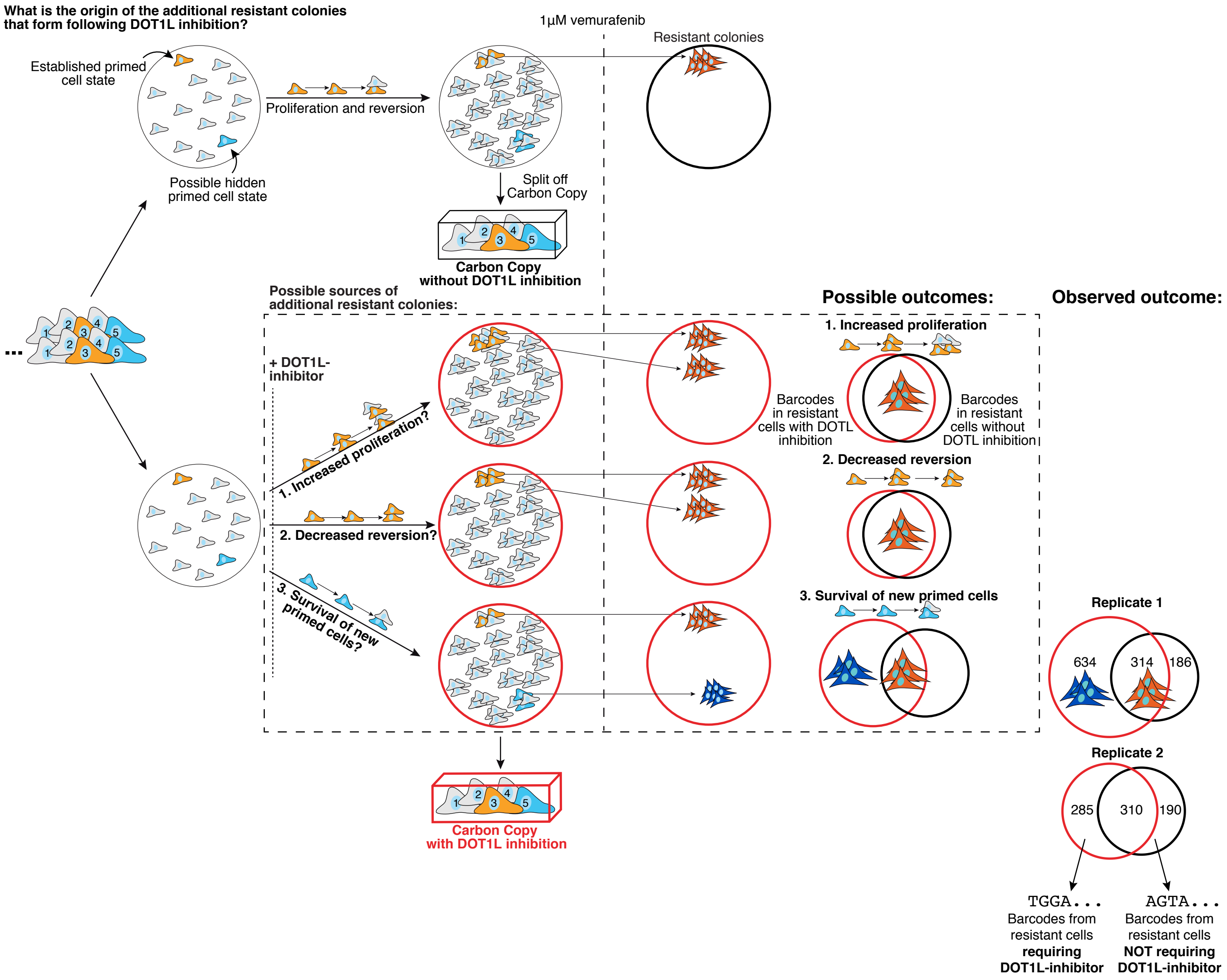


**Supplementary Figure 11. Identification and isolation of cells requiring DOT1L inhibition to become vemurafenib resistant.** WM989 A6-G3 cells transition into and out of a rare primed cell state that gives rise to drug resistant colonies following vemurafenib treatment. We hypothesized that DOT1L inhibition increases the frequency of resistant colonies by either 1. selectively increasing the proliferation of these primed cells, 2. decreasing their transition out of the primed state, or 3. enabling a new subpopulation of cells to survive vemurafenib treatment. We reasoned that we could distinguish these possibilities by splitting barcoded cells into parallel cultures, treating one with DOT1L inhibitor (4 µM pinometostat for 6 days) and the other with vehicle control (DMSO), then treating both with 1 µM vemurafenib and comparing the barcodes in cells that survive. If either 1. increased primed cell proliferation or 2. decreased reversion were the sole factors leading to an increase in vemurafenib resistance, then we expected to find mostly the same barcode sequences in resistant colonies from DOT1L inhibitor and vehicle control pre-treated cultures (with a small number of distinct barcodes similar to what we observed in the "heritability split" experiments; Supplementary Fig. 3). In contrast, if DOT1L inhibition permitted a new subset of cells to survive vemurafenib treatment, then we expected to find their distinct barcodes in the additional resistant colonies that emerge with DOTL inhibitor pre-treatment. Rightmost Venn diagrams show the observed overlap in barcode sequences in resistant cells from cultures pre-treated with DOT1L inhibitor (red) or vehicle control (black). The presence of distinct barcodes in resistant colonies that emerge with DOT1L inhibitor pre-treatment suggests that DOT1L inhibition increases the frequency of resistance, in part, by enabling a new subpopulation of cells to survive and proliferate in vemurafenib. Here, to select barcodes from resistant colonies, we used the normalized read count for the 500th most abundant barcode in the vehicle control pre-treated samples as a threshold. This threshold undercounts the number of resistant colony barcodes in DOT1L inhibitor pre-treated cultures since these cultures have a greater total number of unique barcodes which reduces the relative abundance of each barcode sequence. Therefore, the number of distinct barcodes in vemurafenib resistant colonies forming after DOT1L inhibition is likely greater than what is shown by the Venn diagrams.

#### Supplementary Figure 12

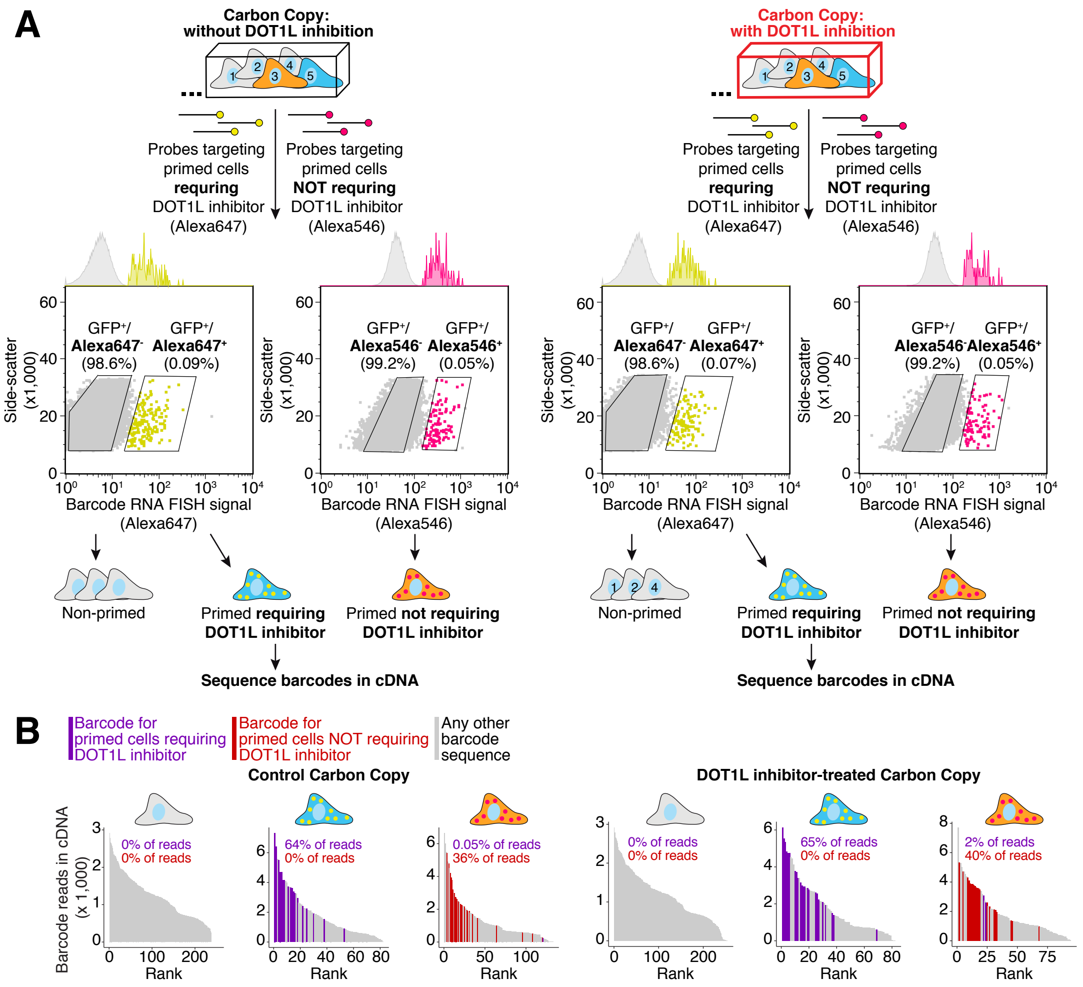


**Supplementary Figure 12. Using Rewind to isolate cells requiring DOT1L inhibition to become vemurafenib resistant and primed cells not requiring DOT1L inhibition to become resistant. A.** As described in Results and Methods, we designed RNA FISH probe sets to distinguish barcode RNA in cells that require DOT1L inhibition to become vemurafenib resistant from barcode RNA in primed cells that do not require DOT1L inhibition to become resistant. FACS plots show the gating used to isolate cells from Carbon Copies labelled with these probe sets (original FCS files and PDFs are available on Dropbox). For our first experimental replicate, we isolated separate cell populations using probes coupled to distinct fluorophores (Alexa Fluor 647 and Alexa Fluor 546). For the second experimental replicate (not shown) we divided our Carbon Copies in two and hybridized each half with separate barcode RNA FISH probe sets coupled to Alexa Fluor 647. We isolated equal numbers of GFP+/Alexa546-/Alexa647- non-primed cells. After sorting, we prepared libraries for RNA sequencing and with the extra cDNA, we performed targeted barcode sequencing. **B.** Bargraphs show the abundance (y-axis) and rank (x-axis) ofbarcodes (≥ 5 normalized reads) sequenced from sorted cells in A. Sequences matching barcodes from cells requiring DOT1L inhibition to become resistant are colored purple. Sequences matching barcodes from cells not requiring DOT1L inhibition to become resistant are colored red. All other barcode sequences are colored gray. Inset shows the percent of barcode sequencing reads that match a probe-targeted barcode.

### Supplementary Figure 13

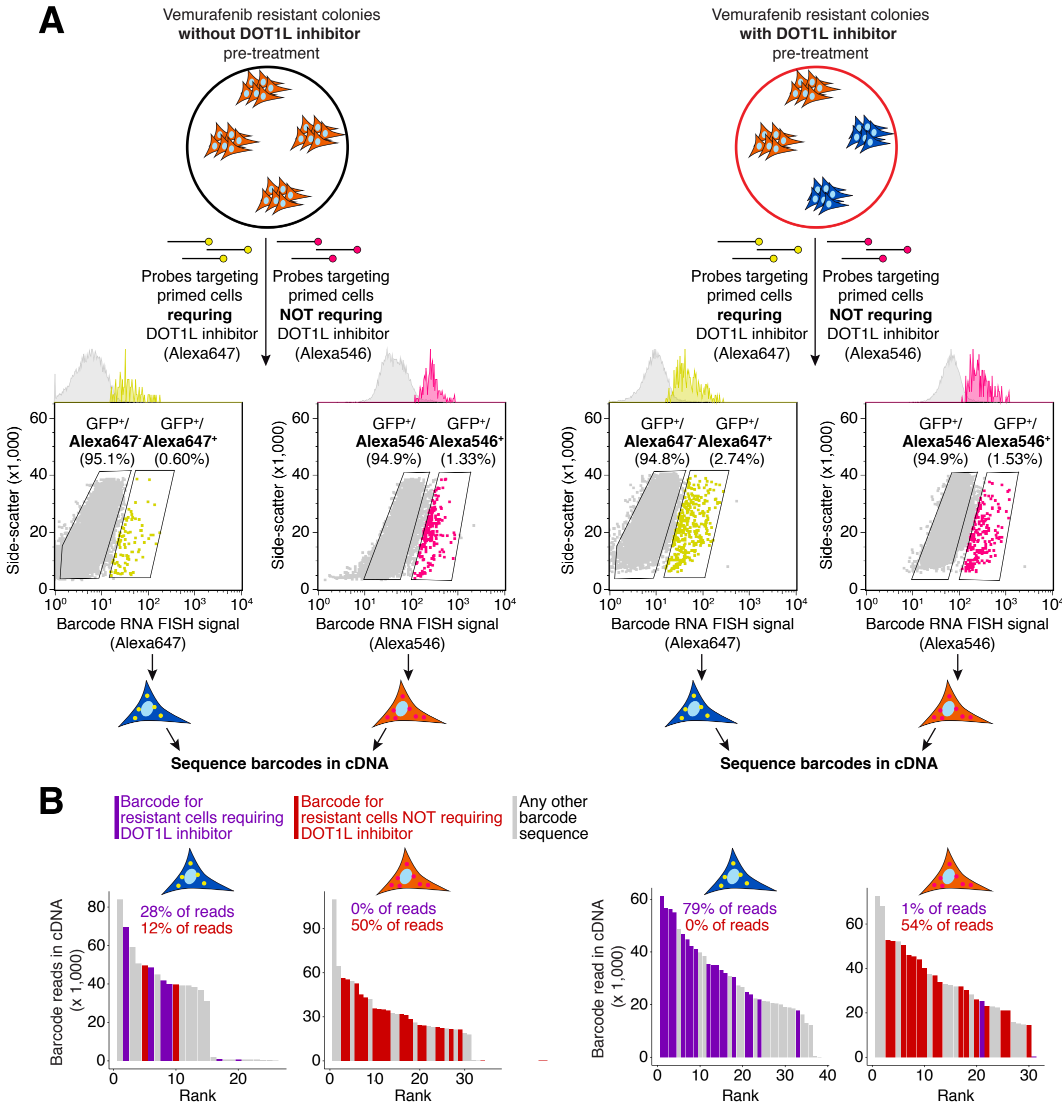


**Supplementary Figure 13. Validation of barcode RNA FISH probe sets targeting cells that require DOT1L inhibition to become vemurafenib resistant and cells that do not require DOT1L inhibition to become resistant. A.** When we performed the Rewind experiments on DOT1L inhibitor-pre-treated and vehicle control-treated cells (see Figure 5 and Supplementary Fig. 9), we fixed 10% of the vemurafenib resistant colonies for validating our barcode RNA FISH probe sets. We expected that probes designed to target cells that require DOT1L inhibition to become resistant to label more resistant cells that were pre-treated with DOT1L inhibitor than resistant cells pre-treated with vehicle control. Conversely, we expected the probes designed to target cells that do not require DOT1L inhibition to label a similar fraction of resistant cells in both conditions. FACS plots show that the probes targeting cells that require DOT1L inhibition labelled approximately 4x as many resistant cells pre-treated with DOT1L inhibitor compared to resistant cells pre-treated with vehicle control, as expected. We observed a minimal difference in labelling using probes designed to target cells that do not require DOT1L inhibition. To verify that the FACS gates correspond to cells expressing the targeted barcodes, we sorted cells from these gates and sequenced their barcodes (original FCS files and PDFs available on Dropbox). **B.** Bargraphs show the abundance (y-axis) and rank (x-axis) for barcodes (≥ 5 normalized reads) sequenced from sorted cells in A. Sequences matching barcodes from cells requiring DOT1L inhibition to become resistant are colored purple. Sequences matching barcodes from cells not requiring DOT1L inhibition to become resistant are colored red. All other barcode sequences are colored gray. Inset shows the percent of barcode sequencing reads that match a probe-targeted barcode.

### Supplementary Figure 14

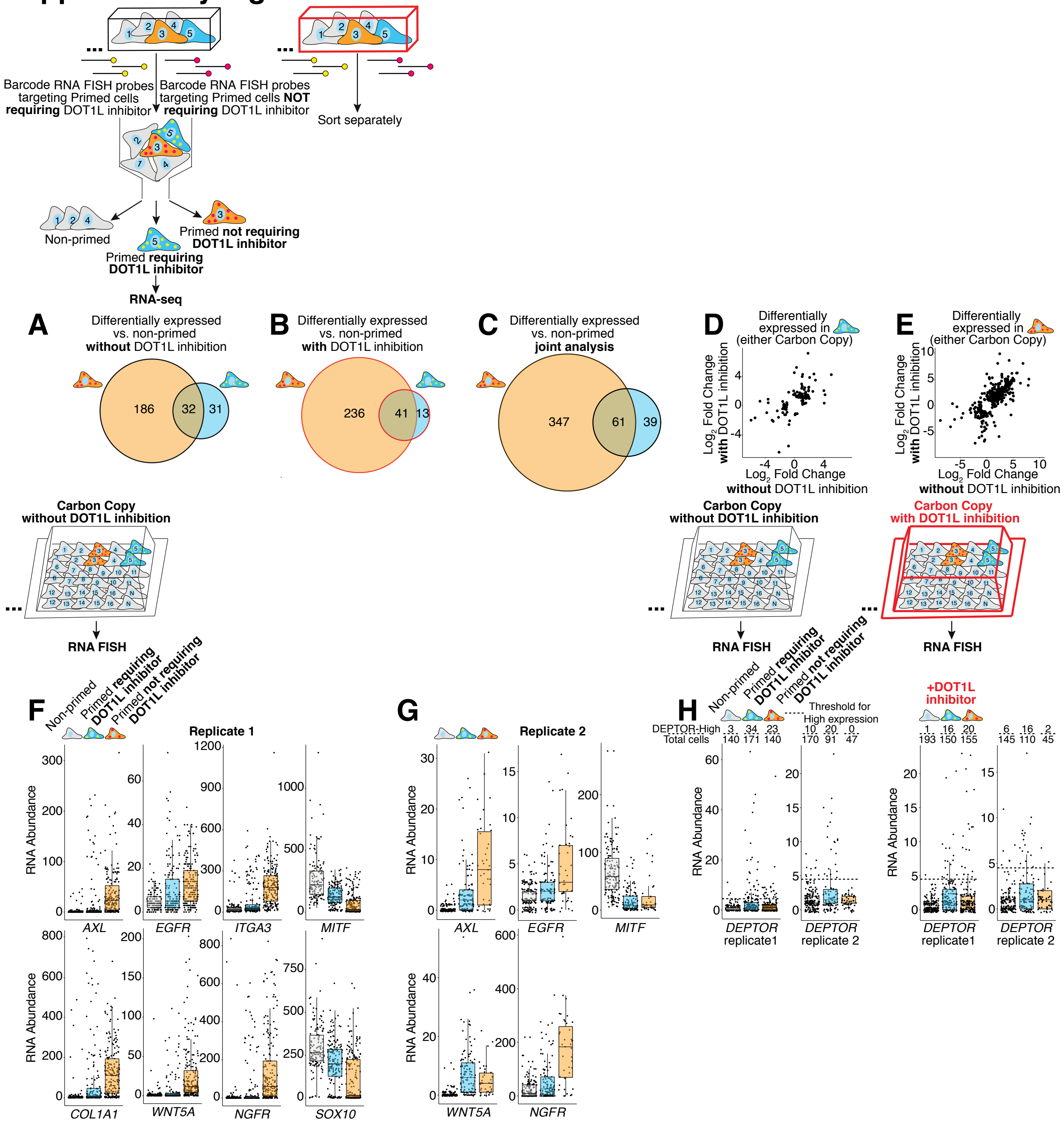


**Supplementary Figure 14. Rewind reveals the transcriptional profile of primed cells that require DOT1L inhibition to become vemurafenib resistant.** We used Rewind to isolate primed cells requiring DOT1L inhibition to become vemurafenib resistant (blue), primed cells not requiring DOT1L inhibition (orange), and non-primed cells (gray). We then measured gene expression in each sorted subpopulation using RNA sequencing. We isolated these subpopulations separately from Carbon Copies treated with DOT1L-inhibitor (red box) or with DMSO (black bock). **A-C.** Venn diagrams show the overlap in differentially-expressed genes (p-adjusted ≤ 0.1 and abs($log_2$ fold change) ≥ 1) identified in each subpopulation of primed cells compared to non-primed cells. A and B show separate analyses for cells isolated from DMSO control and DOT1L-inhibitor treated Carbon Copies, respectively, and C shows the results of a combined analysis with DOT1L-inhibitor treatment modeled as a covariate. Of note, we observed few significant changes in gene expression comparing DOT1L-inhibitor treatment to DMSO within each subpopulation (Supplementary Fig. 16d). **D-E.** For all differentially expressed genes identified in each subpopulation of primed cells, we plotted the $log_2$ fold change with DOT1L-inhibitor treatment (y-axes) vs. with DMSO treatment (x-axes). This analysis revealed that most gene expression differences between primed and non-primed cells exist independently of DOT1L-inhibitor treatment. **F-G.** We used Rewind to identify the indicated subpopulations of cells in Carbon Copies treated with DMSO then measured expression of priming markers in single cells via RNA FISH. Plotted are expression levels in individual cells. Panel F corresponds to the same biological replicate shown in Figure 5. Panel G shows data for a subset of markers measured in a second biological replicate. **H.** Plots show the single-cell expression of *DEPTOR* in the indicated subpopulations of cells measured in Carbon Copies treated with DMSO (left) and Carbon Copies treated with DOT1L inhibitor (right). In both conditions, we found an enrichment of cells expressing high levels of *DEPTOR* (threshold indicated by dotted line; ~ 95th percentile of non-primed cells) in primed cells that require DOT1L inhibition to become vemurafenib resistant. Of note, in 1 replicate we also found an enrichment of cells expressing high levels of *DEPTOR* among primed cells that become resistant independent of DOT1L inhibition. For all boxplots, center line indicates median, box limits indicate 25h and 75th percentiles and whiskers extend to 1.5 times the interquartile range.

### Supplementary Figure 15

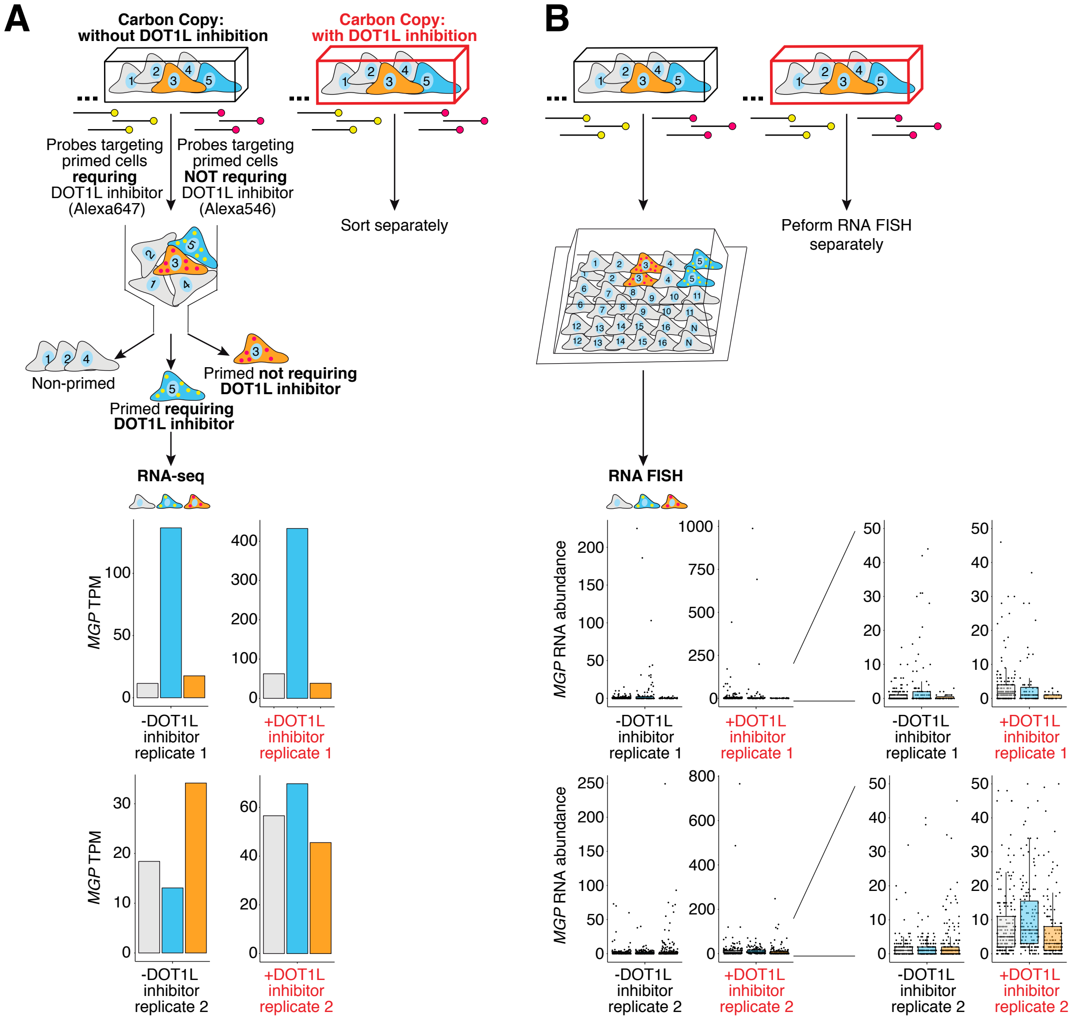


**Supplementary Figure 15. MGP is not a consistent marker of cells that require DOT1L inhibition to become vemurafenib resistant. A.** As described in the Results, we sought to use Rewind to identify markers specific to cells that require DOT1L inhibition to become vemurafenib resistant. In our first experimental replicate, we found that by RNA sequencing primed cells requiring DOT1L inhibition to become resistant (blue) expressed 8-12 fold higher levels of *MGP* compared to either non-primed cells (gray) or primed cells not requiring DOT1L inhibition (orange). This, however, did not replicate in a second experiment. Bargraphs show the *MGP* expression data from these 2 independent biological replicates. **B.** Based on the magnitude of the initial observation, we nonetheless used RNA FISH to compare single-cell expression of *MGP* between subpopulations of cells identified using Rewind in separate sets of Carbon Copies. The first RNA FISH experiment revealed an enrichment of cells expressing high levels of *MGP* in cells that require DOT1L inhibition to become resistant, however this too did not replicate in a second experiment. For all boxplots, center line indicates median, box limits indicate 25h and 75th percentiles, and whiskers extend to 1.5 times the interquartile range.

### Supplementary Figure 16

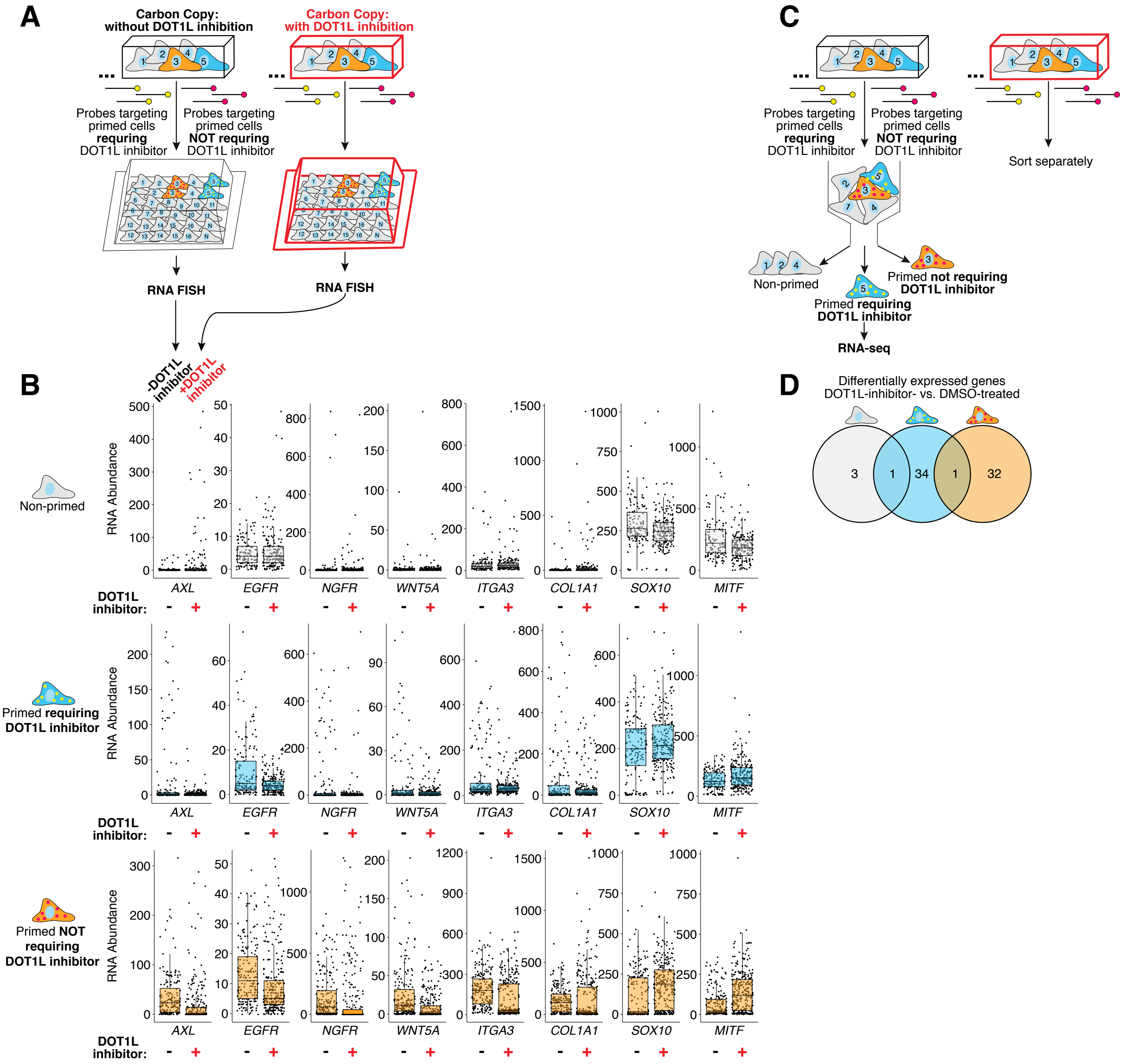


**Supplementary Figure 16. DOT1L inhibition partially decouples expression of established priming markers and vemurafenib resistance A.** As described in the main text, we asked whether DOT1L inhibition enabled a new subpopulation of cells (blue) to survive vemurafenib treatment by converting them into the previously established primed cell state (orange). To address this question, we used Rewind to identify both primed subpopulations (blue and orange) and non-primed cells (gray) in Carbon Copies treated with DOT1L inhibitor (red outline) and the Carbon Copies treated with DMSO (black outline), then used RNA FISH to measure single-cell expression of 8 genes associated with the primed cell state. Six of these genes (*AXL*, *EGFR*, *NGFR*, *WNT5A*, *ITGA3*, and *COL1A1*) have higher expression in the primed cell state while *SOX10* and *MITF* have lower expression in the primed cell state. **B.** We found that DOT1L inhibition modestly increased expression of several genes elevated in the established primed cell state (*AXL*, *NGFR*, *COL1A1*) and decreased expression of *SOX10* and *MITF* specifically in non-primed cells that do not ultimately survive vemurafenib treatment (top row, gray). In contrast, for cells that do ultimately survive vemurafenib treatment (blue and orange), DOT1L inhibition appeared to decrease expression of positive markers of the established primed cell state and increase expression of *SOX10* and *MITF*. These transcriptional changes, away from the established primed cell state, may suggest that compared with vemurafenib treatment alone, cells pre-treated with a DOT1L inhibitor can become vemurafenib resistant via an alternate path. For all boxplots, center line indicates median, box limits indicate 25h and 75th percentiles and whiskers extend to 1.5 times the interquartile range. **C-D.** As in Figure 5, we performed bulk RNA sequencing on each subpopulation of cells sorted separately from the Carbon Copies treated with DOT1L inhibitor and the Carbon Copies treated with DMSO. For each subpopulation, we identified differentially-expressed genes (p-adjusted ≤ 0.1 and abs($log_2$ fold change) ≥ 1) comparing cells treated with DOT1L inhibitor vs. DMSO, then plotted the overlap across subpopulations. Notably, the number and magnitude of signficant gene expression changes induced by DOT1L inhibition **within** each subpopulation is much lower than the gene expression differences observed between subpopulations (Supplementary Fig. 14).
